# The value of what’s to come: neural mechanisms coupling prediction error and reward anticipation

**DOI:** 10.1101/588699

**Authors:** Kiyohito Iigaya, Tobias U. Hauser, Zeb Kurth-Nelson, John P. O’Doherty, Peter Dayan, Raymond J. Dolan

**Affiliations:** Max-Planck UCL Centre for Computational Psychiatry and Ageing Research, 10-12 Russell Square, London WC1B 5EH, United Kingdom; Gatsby Computational Neuroscience Unit, University College London, 25 Howland Street, London, W1T 4JG, United Kingdom; Division of Humanities and Social Sciences, California Institute of Technology, 1200 E California Blvd, Pasadena, CA 91125; Wellcome Centre for Human Neuroimaging, University College London, 12 Queen Square, London WC1N 3BG, United Kingdom; Deepmind, 6 Pancras Square, London N1C 4AG, United Kingdom; Max Planck Institute for Biological Cybernetics, 72076 Tubingen, Germany

## Abstract

Having something to look forward to is a keystone of well-being. Anticipation of a future reward, like an upcoming vacation, can often be more gratifying than the very experience itself. Theories of anticipation have described how it induces behaviors ranging from beneficial information-seeking through to harmful addiction. However, it remains unclear how neural systems compute an attractive value from anticipation, instead of from the reward itself. To address this gap, we administered a decision-making task to human participants that allowed us to analyze brain activity during receipt of information predictive of future pleasant outcomes. Using a computational model of anticipatory value that captures participants’ decisions, we show that an anticipatory value signal is orchestrated by influences from three brain regions. Ventromedial prefrontal cortex (vmPFC) tracks the value of anticipation; dopaminergic midbrain responds to information that enhances anticipation, while sustained hippocampal activity provides a functional coupling between these regions. This coordinating function of the hippocampus is consistent with its known role in episodic future thinking. Our findings shed new light on the neural underpinnings of anticipation’s influence over decision-making, while also unifying a range of phenomena associated with risk and time-delay preference.

## Introduction

*“Pleasure not known beforehand is half-wasted; to anticipate it is to double it.”* – Thomas Hardy, *The Return of the Native*

Standard economic theory suggests a reward is more attractive when it is imminent (e.g. eating now) than when it is delayed (e.g. eating tomorrow), predicting that people always consume a reward immediately. This so-called temporal discounting^1–3^ has been adapted with great success, such as in the design of artificial intelligence systems that can plan their future effectively^4–6^ through to understanding aspects of the human mind.^7–10^ However, real life behavior is more complex.^11–15^ Humans and other animals sometimes prefer deliberately to postpone pleasant experiences (e.g. saving a piece of cake for tomorrow, or delaying a one-time opportunity to kiss a celebrity^11^), clearly contradicting simple temporal discounting.

An alternative idea in behavioral economics poses that we enjoy, or *savor*, the moments leading up to reward. This idea, formally known as the utility of anticipation,^11, 13, 16–19^ entails that people experience positive (utility) value while waiting for a reward, in addition to the value they derive from its ultimate consumption. The added value of anticipation naturally explains why people occasionally prefer to delay reward (e.g. because we can enjoy the anticipation of eating a cake until tomorrow by saving it now),^11^ as well as a host of other human behaviors such as information-seeking and addiction.^16, 20^

However, with notable exceptions,^19^ few studies have examined the neural basis of the value arising from reward anticipation. This contrasts with ample studies examining the value arising from reward itself^21–24^ and its temporal discounting^7, 10, 25–27^ (which are sometimes confusingly referred to as reward expectation; however, the values do not arise from anticipation but from reward itself). Further, no studies have linked neural responses during anticipatory periods^19, 28–31^ to anticipatory values that drive subject’s actual decisions. Therefore, despite the theory’s behavioral explanatory power, there is a major gap in understanding the neural roots of anticipatory values.

Here we investigated the neurobiological underpinnings of value computation arising from reward anticipation, by combining computational modeling, a behavioral task, and functional magnetic resonance imaging (fMRI). We fit our computational model^20^ of anticipatory value^11^ to task behavior, and for each participant use best model to make predictions about the timecourse of anticipatory value in the brain for each participant. We then compared this predicted signal with actual fMRI data, finding that the ventromedial prefrontal cortex (vmPFC) encoded the temporal dynamics of an anticipatory value signal and dopaminergic midbrain encoded a signal reporting changes in reward expectation. This reward prediction error (RPE) is widely interpreted as a learning signal in reinforcement learning theory,^23, 32–34^ but our model predicts it can also act to enhance anticipatory value, which in turn drives behavior. We show that hippocampus mediates this enhancement of value, being associated with a functional coupling between the other two regions (the vmPFC and the dopaminergic midbrain).^35–41^ In light of a strong tie between the hippocampus and memory and future imagination, we conclude our data are consistent with a notable suggestion from behavioral economics that the value of anticipation relates to a vivid imagination of future reward.^42–48^

## Results

### Participants prefer to receive advance information about upcoming reward

We employed the same behavioral task that has previously been linked to the value of anticipation (**Figure 1a**). In brief, our task examines how participants change their preference for resolving uncertainty about future pleasurable outcomes, according to reward probability and delay duration until outcomes (please also see the Methods section). Participants made decisions with full knowledge regarding conditions (probability, and delay, of reward outcomes) as these were signaled with simple visual stimuli on each trial. The conditions were randomly selected for each trial — the probability was sampled uniformly at random from 0.05, 0.25, 0.5, 0.75, 0.95, and the duration of a wait period until reward or no-reward delivery was sampled uniformly at random from 1, 5, 10, 20, 40 s.

**Figure 1:**
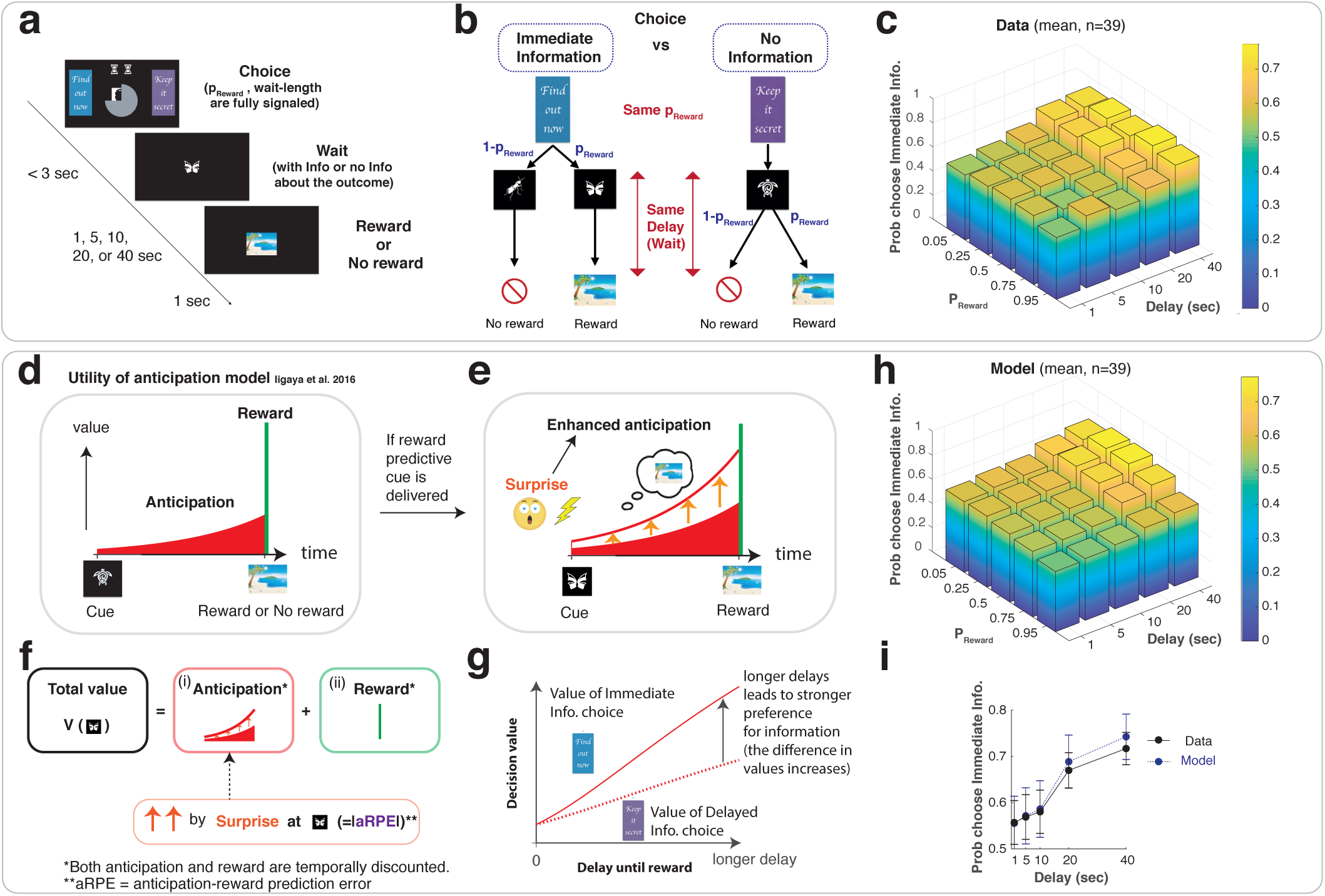
The value of anticipation drives a preference for advance information. (**a**). Task. On each trial, participants were presented with two lateral choice targets: an immediate-information target (‘Find out now’) and a no-information target (‘Keep it secret’) as well as two central stimuli signaling the probability of reward (sampled uniformly at random from 0.05, 0.25, 0.5, 0.75, 0.95) and the duration of a wait period until reward or no-reward delivery (sampled uniformly at random from 1, 5, 10, 20, 40 s). Once participants chose a target, a symbolic image cue was presented and remained present for the entire wait period. After this period, a rewarding image or an image signaling no reward appeared. (**b**). The immediate-information target was followed by symbolic cues that predict upcoming reward or no reward with no remaining ambiguity (reward predictive cue, or no-reward predictive cue). The no-information target was followed by a cue that implied nothing about the reward outcome (no-information cue), so that the future outcome remained uncertain during this wait period. (**c**). Average behavior. Participants showed a stronger preference for advance information under longer delay conditions. The effect of reward probability was not significant as a group (two-way ANOVA, *F*_4, 950_ = 10.0, *p* < 0.001 and *F*_4, 950_ = 0.35, *p >* 0.05, respectively), but showed heterogeneous dependencies (**Figure S4**), consistent with our model. (**d**,**e**,**f**,**g**). Computational model.^20^ (**d**). Following a standard characterization of the utility of anticipation,^11^ our model assumes that the value of each cue is determined by the sum of (i) the value of anticipation that can be consumed while waiting for reward (red), and (ii) the value of reward consumption itself (green). (**e**,**f**). If the immediate-information target is chosen, and a reward predictive cue is presented, the value of anticipation is boosted throughout the delay period (orange upward arrows). The boosting is quantified by a surprise, proportional to the absolute value of prediction error that is computed from both the value of anticipation and reward (anticipation-reward prediction error: aRPE, Eq. 1). (**g**). The model predicts that the value difference between the immediate-information target and the no-information target is larger under longer delay conditions, due to the sustained boosting.^20^ (**h**). The average of modeled preferences across participants. We fit our model to participants’ behavior using a hierarchical Bayesian procedure.^20, 49^ (**i**). The model (blue) captures the effect of delay conditions in data (black). The error bars indicate the mean and standard errors of participants (*n* = 39). The model also captures the effect of probability conditions (**Figure S2**). See **Figure S1** for how other classical models fail to capture the data.

On each trial, participants chose between an immediate-information target (labeled ‘Find out now’) and a no-information target (‘Keep it secret’). If the immediate-information target was chosen, one of two cues, each of which uniquely signaled if reward would or would not arrive, was shown during the wait period (**Figure 1b**, left). If the no-information target was chosen, a separate non-predictive cue that carries no information about an upcoming outcome was shown on the screen during the wait period (**Figure 1b**, right), ultimately followed by either reward or no-reward. The reward image was randomly drawn from previously validated rewarding pictures,^20, 50^ and consequently subject to immediate consumption (by viewing) upon delivery. The no-reward image was a neutral image indicating no-reward.

In this design, participants’ choices did not affect final reward outcomes or duration of delays (**Figure 1b**). Both reward probability and delay duration were pre-determined and signaled to participants at the beginning of each trial. Participants could only choose if they want to gain knowledge about whether they would receive a reward or not before a delay. Therefore, standard decision theories that aim to maximize the chance of receiving rewards would predict no preference over two choices, because the probability of obtaining a reward (hence the expected value) is the same across the two choices (please see **Figure S1abc**). Thus models with conventional temporal-discounting predict no preference.

However, against the predictions of conventional theory, we found that participants exhibited a preference for advance information. Further, consistent with previous findings,^51–53^ the preference for immediate information increased with delay^20, 54^ (**Figure 1ci**).

### A computational model of anticipatory values accounts for participants’ choice

We accounted for the preference for obtaining advance information using an economic notion of the utility of anticipation.^11, 13, 16–19, 55^ While standard value-based decision theories assign values to consummation of reward itself, theories of the utility of anticipation also assign values to the moments up until receipt of reward (**Figure 1d**; see Eq.1 in the Methods section). One possible psychological root for the value arising from reward anticipation is an enjoyable subjective feeling while waiting for pleasant outcomes;^11^ but the mathematical framework of the anticipatory value is open to wider interpretations (e.g.^16, 56^). Here we examine how the mathematical model can account for choice behavior and test its predictions on neuroimaging data. We will discuss psychological interpretations following this.

Although the value of anticipation naturally accounts for why people want occasionally delay rewards (because they can consume anticipatory utility while waiting), the original formulation does not necessarily explain a preference for obtaining advance information. The model still predicts indifference between the two choices in the task, because the value of anticipation is linearly scaled with the probability of reward (as is the case for the expected value of the actual outcome), leading to the same average values for two choices^20^ (illustrated in **Figure S1def**). This is expected because information plays little role in the original formulation.

Therefore, we recently proposed a slight modification to this original formulation.^20^ The idea here is that people receive *enhanced* value of anticipation when they unexpectedly discover a reward is impending (**Figure 1ef**). This surprise-enhancement of anticipatory value is inspired by experimental observations that such unexpected discoveries lead animals to become, and remain, more excited than when animals waited for a certain reward with no such surprising discovery,^54^ where animals paradoxically prefer a less rewarding but more surprising choice.^57^

We mathematically formulated a surprise that relates to the enhancement of anticipatory value by using a notion of reward prediction error. Every time participants received advance information about future reward (or no-reward), participants experienced a reward prediction error, defined by the difference between (a) the value of future that is just updated thanks to the arrival of new information and (b) the value of future that was expected before the arrival of information.^58^ While in a standard theory reward prediction error (RPE) is computed from the value of reward, in our model it is computed from the value of anticipation and reward (Eq. 5). Therefore, we refer to our model’s prediction error signal as anticipation+reward prediction error (aRPE) signal.

In our computational model, this aRPE quantifies a surprise that links to enhancement (boosting) of anticipatory value. Following the conventional mapping of prediction error to surprise,^59^ the model quantifies surprise by the absolute value of the aRPE, because unexpected negative outcomes (negative aRPE) can be just as surprising as unexpected positive outcomes (positive aRPE). This also avoids unreasonable effects such as turning negative anticipation to positive anticipation by multiplying by a negative aRPE. A simplest expression for boosting is thus to assume that anticipatory value is linearly enhanced by the absolute value of aRPE (please see Eqs. 1 and 2 in the Methods section). It is important to note that an aRPE (or a standard RPE) is expected to be a phasic signal that lasts only for a short period. However, animals appear to remain excited over the course of whole anticipatory periods,^54^ and so in the model, the enhancement of anticipation is sustained throughout a wait period^20^ (Eqs. 1 and 2). Therefore, the model predicts a signal that is associated with boosting anticipatory values to be a prolonged representation of the absolute value of aRPE (or a prolonged signal that is proportional to the amount of surprise). Such a signal is likely to be encoded in regions other than those encoding phasic aRPEs. We return to this question later in the Results section.

In our task, the cue predictive of a future outcome that follows the immediate-information target creates a dopaminergic aRPE, and it triggers a boosting of the value of anticipation. On the other hand, the non-predictive cue following the no-information target does not generate aRPE, thus it does not trigger any boosting (**Figure S1ghi**). Therefore, the model predicts that participants experience enhanced anticipatory value after receiving a reward predictive cue following the immediate-information target, while they experience a default amount of anticipatory values weighted by probability of reward after receiving a no-information cue following the no-information target. Due to the sustained boosting, the model predicts that the difference in the values between the immediate-information target and the no-information target is larger under longer delay conditions (at least in the absence of strong discounting), causing a stronger preference for the immediate information target at longer delay conditions (**Figure 1g**).^20^

We fit this model to participants’ trial-by-trial behavioral data using a hierarchical Bayesian scheme^20, 49^ (see Methods section). As before,^20^ the model captured participants’ preferences for advance information (**Figure 1hi**). In particular, the model quantitatively captured the key feature of the data, which is the increase in preference for immediate information under longer delay conditions (**Figure 1i**), as well as the preference over probability conditions (**Figure S2**).

Other standard models cannot capture a preference for advance information. For example, models with discounted reward but with no anticipatory value, or models with both discounted reward and anticipatory value but no enhancement of anticipation, cannot capture behavior (please see **Figure S1** for illustration). We formally tested this by fitting other possible models to the behavioral data, and compared the models’ integrated Bayes Information Criterion scores (iBIC;^20, 49, 60^ please see the Methods section), which strongly favored our full model over other standard computational models (**Figure S3**).

In addition to the task behavior here, our model also captures a wide range of existing findings about information-seeking behavior, and offers potential links to addiction and gambling^20^ (also see the Discussion). However, despite the behavioral explanatory power, the neural roots of anticipatory value and its boosting are largely unknown. Therefore, we next sought to elucidate the neurobiological basis of value arising from anticipation, using our computational model that captures participants’ behavior. In particular, we aimed to pin down the neural basis of three key components of our model: namely, the representation of anticipatory value during wait periods; the aRPE signal at advance information cues, and the sustained boosting signal of anticipation during wait periods following surprise. Finally, we examined how brain regions encoding these computational components are coupled together to dynamically orchestrate the value of anticipation.

Here we stress to readers that our computational model is a very general mathematical model, which does not specify the psychological roots of anticipatory value, analogously to standard reinforcement learning models encompassing very complex psychological roots of reward value.^58, 61^ Thus, our goal was to elucidate neural correlates of our computational model’s mathematical predictions about values arising during anticipatory periods, which in turn drive behavior. We discuss possible psychological roots of anticipatory values in the Discussion section.

### The vmPFC encodes our computational model’s value of anticipation

Our model predicts that the value of anticipatory utility dynamically changes over the course of a delay period (Eq.22 in the Methods section). Regardless of boosting, the signal ramps up as the outcome approaches, but the value is also subject to conventional discounting. This implies a tilted inverted-U shape over time under typical parameter settings (**Figure 1e**).

Based on our hierarchical model fit to choice behavior, we calculated each subject’s maximum a posteriori (MAP) parameters within the computational model. Using these parameters, we estimated subject-specific time courses of several variables that we tested on the neural data. The predictions include (i) anticipatory positive and negative value during wait periods (Eq. 22 in the Methods section) (ii) discounted reward value (standard expected reward) during the same periods (Eq. 24 in the Methods section), and (iii) prediction errors at cue presentation (Eq. 28 in the Methods section). These signals were convolved with SPM’s default canonical HRF (**Figure 1h**; see **Figure S5** for an example). As illustrated in the Methods section, we separated predictive anticipatory signals for positive reward and no reward, because we found that participants assigned a negative value to no-reward outcome.^20^

We found the model’s anticipatory value signal for positive reward correlated significantly with BOLD in ventromedial prefrontal cortex (vmPFC) (*p* < 0.05, whole brain FWE correction; peak MNI coordinates [10, 50, 16], *t* = 6.02; **Figure 2a**), as well as dorsal caudate (*p* < 0.05, whole brain FWE correction; peak coordinates [−20, −2, 18], *t* = 5.81 **Figure S6**). These results are consistent with the representation of the value of imagined reward reported previously in vmPFC,^63^ and of reported anticipatory activity in vmPFC^19, 31, 64^ as well as in caudate.^28, 65–68^ Across the brain, we found no significant effect of anticipatory utility arising from no-reward outcome that survived a stringent whole-brain correction (see **Figure S7**). Thus, we focus on the anticipatory value of future reward referred to henceforth as anticipatory value.

**Figure 2:**
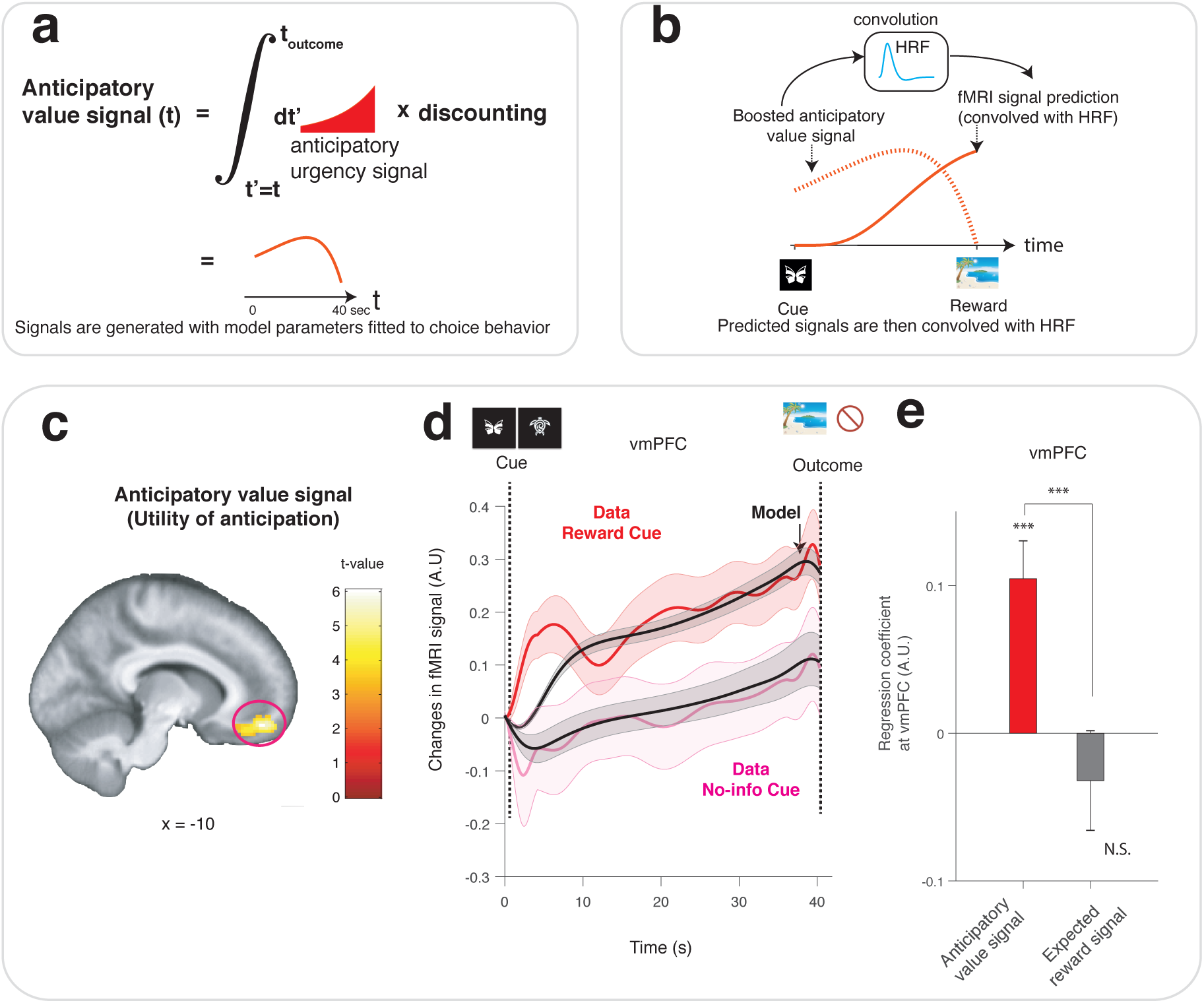
Neural representation of our computational model’s anticipatory value signal in the vmPFC. (**a**). The model’s anticipatory value signal at time *t* is an integral of discounted future anticipation (urgency signal) at *t*^*t*^ *> t*. This integrated signal evolves dynamically during the waiting period (red curve). Note that this signal is different from a well-studied expected value signal of future reward, which is also included as a separate regressor in the same GLM analysis. (**b**). The model’s prediction for fMRI signals (solid red) is computed by convolving the value signal (dotted red) with the canonical haemodynamic response function (light blue). (**c**). BOLD in vmPFC positively correlated with an anticipatory value signal. This slow correlation survived our phase-randomization test that is protected for whole brain FWE (*p* < 0.001; see **Figure S8**), as well as SPM’s standard whole brain FWE correction (*p* < 0.05). A cluster of voxels with *p* < 0.001 (uncorrected) surrounding the peak [10,50,16] is shown for display purposes. Please also see **Figure S6** for other regions, including caudate. (**d**). The temporal dynamics of the BOLD signal in the vmPFC cluster (shown in Figure 2c) matched the model’s anticipatory value signal during the anticipation period. Changes in activity averaged over participants following receipt of a reward predictive cue (Red), and after receipt of a no-information cue (magenta), are shown, as well as the model’s prediction for each of these conditions (black). The error bar indicates the SEM over participants. The data was upsampled for this plot, following.^62^ (**e**). A control, confirmatory, analysis shows that the vmPFC is more strongly correlated with our model’s anticipatory utility signal than a standard expected (discounted) reward value signal. In the GLM analysis with both regressors, average regression weights in the vmPFC cluster for the anticipatory value signal were significantly greater than the coefficients to the expected reward signal (*p* < 0.001, permutation test). The average regression weights in the vmPFC cluster were significantly larger than zero for our model’s predicted signal (*p* < 0.001 t-test, *t*_38_ = 4.07), but not significantly different from zero for the expected reward signal. The error bars indicate the mean and SEM. Note that this is a confirmatory analysis. See **Figure S9** for correlates with the expected reward expectation signal.

Given the need to avoid potential false positives from auto-correlations in slowly-changing signals,^69^ we conducted non-parametric, phase-randomization, tests wherein we scrambled the phases of signals in a Fourier decomposition^70–72^ (**Figure S8a**). This test can be generally applied to neuroimaging and electrophysiology studies, so as to avoid false positive discoveries particularly when analyzing correlations between slow signals such as values.^69, 71^To do so, we transformed our model’s predicted anticipatory value signal for each participant into Fourier space, randomized the phase of each frequency component, and transformed the signal back to the original space. Only the regressor being tested was randomized, while others were kept the same in the full GLM. We then performed a standard analysis on this full GLM for each participant with the scrambled signal, and then conducted a second level analysis. By repeating this procedure many times, we created a null distribution. To protect this test against family-wise error, we constructed the null distribution by taking a maximum value of correlation score across a region of interest, or across the whole brain, from each of our second level analyses, comparing against the correlation value in the original analysis. We found that the effects in the vmPFC (*p* < 0.001 randomization whole-brain FWE corrected) and the caudate (*p* < 0.01 randomization whole brain FWE corrected) survived this Fourier phase randomization test (**Figure S8b**).

A more detailed inspection of these signals, during the wait period, showed the time course of vmPFC activity closely resembled our model’s predictions. In **Figure 2b**, we plot the time course of average fMRI signals in the vmPFC cluster shown in **Figure 2a** during the wait period separately for two conditions, namely when participants received a reward predictive cue (red), and when participants received a no-information cue (magenta). The time-courses track the model’s predictions in each condition (black).

We note that the vmPFC cluster was more strongly correlated with our model’s anticipatory value signal than with a standard expected reward signal (our discounted reward value signal) that was also present in the same GLM, though both of the signals showed a similar ramping toward reward (please see **Figure S5** for an example participant). As a confirmatory analysis, in **Figure 2e** we plotted average beta values in the vmPFC cluster for the anticipatory value and for the standard expected reward (note: both regressors are present in the same GLM) and confirmed that the difference between the coefficients was significant (*p* < 0.001 permutation test). The model’s expected reward signal was instead correlated significantly with regions including the superior temporal gyrus (*p* < 0.05, whole brain FWE correction; [−48, −48, 16], *t* = 5.28, **Figure S9a**). This also survived a phase-randomization test (*p* < 0.001).

We further asked whether BOLD in the vmPFC during the wait period correlated with a simpler signal, such as constant expected outcome value. When the immediate-information cue is presented, this is the same as the value of reward or no-reward without discounting or anticipatory modulation; otherwise, it is an average of the values of reward and no-reward weighted by their respective probabilities. We examined the singular contribution of this signal by adding it as another parametric boxcar regressor during waiting periods to the original GLM, and then compared the average beta values of the vmPFC cluster between the anticipatory value, and the expected value, regressor. In this way, we estimated the partial correlation of each regressor. As shown in **Figure S10**, the vmPFC BOLD was more strongly correlated with the model’s anticipatory value signal than with the constant expected value signal (*p* < 0.001 permutation test). BOLD was still positively correlated with the model’s anticipatory value signal (*p* < 0.001 t-test, *t*_38_ = 3.93), and the effect of an expected value signal was not significant.

For completeness, we report descriptively that an anticipatory urgency signal, which is an anticipation signal before integration (Eq. 26 in the Methods section) correlated with anterior insular cortex^29^ ([34, 30, 2], phase-randomization test *p* < 0.01, **Figure S9b**).

### The dopaminergic midbrain encodes our computational model’s anticipation-reward prediction errors (aRPE) at the time of advance information cues

The anticipation-reward prediction error (aRPE) arising at advance information cues is a unique and critical signal in our model. First, unlike conventional models relying on reward, our model’s aRPE is computed from the value arising from both reward anticipation and reward itself (Eq. 5 in the Methods section). Second, while in a standard reinforcement learning model an RPE serves as a learning signal, in our model it triggers a surprise that is associated with enhancement (boosting) of anticipatory value (Eq. 2 in the Method section).

In this regard aRPE also differs from a conventional temporal difference (TD) prediction error,^4, 32^ which only considers conventionally discounted outcomes and does not involve boosting. Rather, our computational model’s aRPE signal encompass both a standard RPE and the so-called information prediction error (IPE),^52, 53, 74, 75^ both of which have been previously shown to be represented in the activity of dopamine neurons.^23, 32–34, 52^ Dopamine has been also implicated in enhanced motivation. ^76–79^

Therefore, based on extensive prior studies, we hypothesized that an aRPE signal arising at the time of advance information cues would be encoded in the midbrain dopaminergic regions and the ventral striatum (e.g.^23, 30, 52, 73, 80^). For this, using each subject’s MAP parameter estimates obtained from fitting our model to choice behavior, we calculated a full, signed, aRPE signal, occasioned at the onset of advance information cues (reward predictive, no-reward predictive, and no-information cues), based on the discounted value of anticipation (including both positive and negative cases) and that of outcomes (Eq. 28).

We assumed that participants fully learned the task in the training period. Therefore, the size of aRPE was fully determined by each trial’s experimental conditions (probability and delay of reward) as well as the model’s fitted parameters, meaning that an aRPE was not affected by recent trials’ outcomes. Therefore, we analyzed the fully self-consistent aRPE (Eq. 28 in the Method section).

We found that the model’s signal correlated significantly with BOLD in a midbrain dopaminergic region encompassing the ventral tegmental area and substantia nigra (VTA/SN) (**Figure 3a**; *p* < 0.05, small volume FWE correction with an anatomical ROI;^73^ [4, −26, 20], *t* = 3.78). Additionally, we found also that BOLD in the medial posterior parietal cortex (mPPC)^81, 82^ correlated significantly with the model’s predicted signal (**Figure 3a**; *p* < 0.05, cluster-level whole brain FWE correction with the height threshold *p* < 0.001; *k* = 166, peak at [0, −42, 50].). We did not find significant associations in the ventral striatum, perhaps because cue- and reward onsets were unusually temporally distant (up to 40 sec), a finding consistent with a previous report that the anticipation of reward in a music piece triggers a dopamine release in the caudate, but not in the ventral striatum.^67^

**Figure 3:**
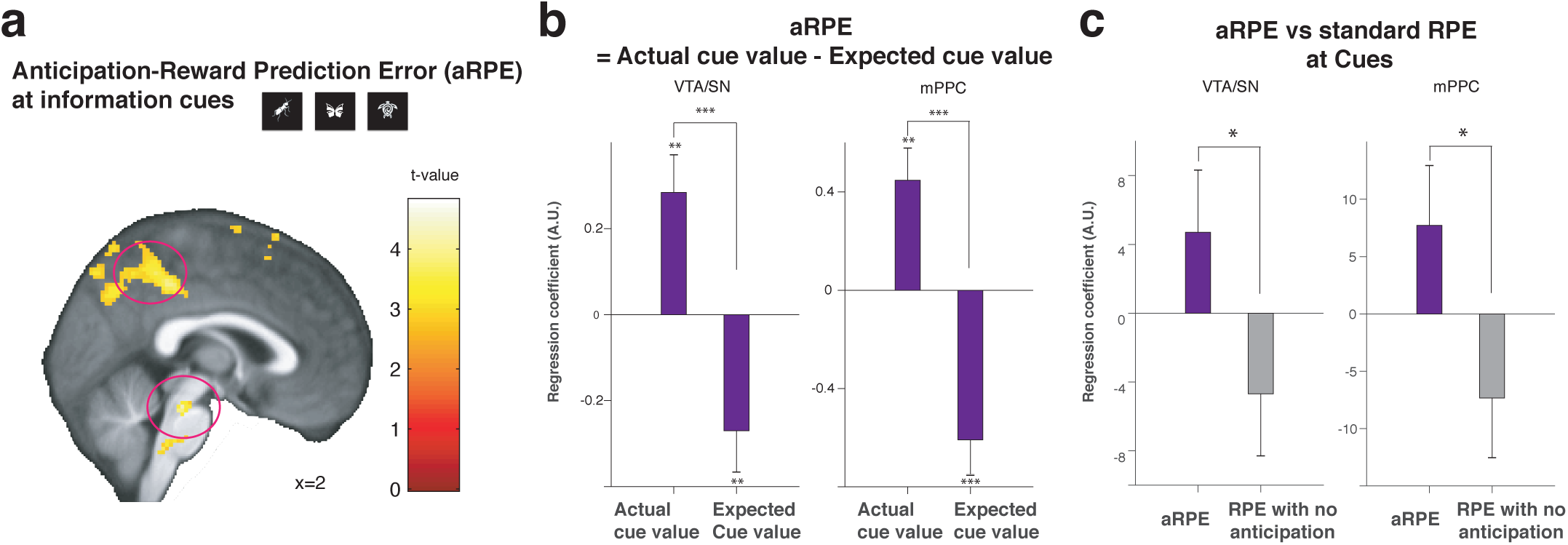
Neural representation of our computational model’s anticipation-reward prediction error (aRPE) signals at information cues. (**a**) The VTA/SN and mPPC BOLD positively correlated with the model’s aRPE at the time of advance information cue presentations (VTA/SN: *p* < 0.05 FWE small volume correction,^73^ mPPC: *p* < 0.05 whole-brain FWE, cluster corrected at *p* < 0.001). All three cues (reward predictive, no-reward predictive, and no-information cues) were included as onsets. Voxels at *p* < 0.005 (uncorrected) are highlighted for display purposes. The model’s aRPE was computed from the values of anticipation and reward (Eq. 28 in the Methods section), in contrast to standard RPE signals in reinforcement learning models. (**b**) Our confirmatory analysis shows both the VTA/SN and the mPPC show paradigmatic correlations with aRPE. At the time of advance information cue presentations, BOLD in the VTA/SN and the mPPC positively correlated with the model’s actual cue value signal, and negatively with the model’s expected cue value signal, indicating both regions express canonical prediction errors. The difference of the mean coefficients was significant in the VTA/SN (*p* < 0.001 permutation test) and in the mPPC (*p* < 0.001 permutation test). The positive correlation with cue outcome values and the negative correlation with expected values were all significant (received cue value: *p* < 0.01 for the VTA/SN and the mPPC by t-test *t*_38_ = 3.24 and *t*_38_ = 3.40, expected cue value: *p* < 0.01 for the VTA/SN and *p* < 0.001 for the mPPC by t-test *t*_38_ = 2.82 and *t*_38_ = 4.37). The two regressors were included in the same GLM. (**c**). Our confirmatory analysis shows both regions express stronger correlations with our model’s full aRPE than with standard prediction error with discounted reward (RPE) at advance information cues. The difference was significant between the mean coefficients in the VTA/SN and in the mPPC cluster (*p* < 0.05 permutation test). The two regressors were included in the same GLM. The three stars indicate *p* < 0.001, two stars indicate *p* < 0.01, and one star indicates *p* < 0.05.

Previous studies suggest that significant correlations reported between fMRI signals and prediction errors might be attributable to strong correlations with actual cue value alone, regardless of the presence of negative correlations with expected cue value.^62, 83, 84^ To rule out this possibility, we performed a confirmatory analysis by constructing a GLM with separate regressors for the model’s values of presented cue values and the model’s expected cue values, both of which were computed from the value of anticipation and reward (Eq. 5). The average regression coefficients positively correlated with model’s (actually presented) cue value and negatively correlated with model’s expected (average) cue value (**Figure 3b** in both the VTA/SN and in the mPPC clusters shown in **Figure 3a**. Thus, responses in these regions had characteristic of canonical prediction error signals.^62, 83, 84^

Because our model’s aRPE signal, with the values of anticipation and reward, is more complex than a standard RPE signal with reward value alone, we then performed another confirmatory analysis. Here we constructed a GLM that included the model’s full aRPE signal (Eq. 8) and a standard RPE error signal based exclusively on reward values (Eq. 30), and then compared the partial correlations associated with these regressors. We found in both VTA/SN^73^ and the mPPC cluster that the average partial correlation is greater for our model’s full aRPE signal than for the standard RPE signal with discounted reward value alone (**Figure 3c**).

Finally, BOLD in the mPPC has previously been reported to covary with a simpler prediction error signal, the state prediction error signal.^85^ In our experiment, this state-prediction-error (SPE) signal is the absolute value of the difference between outcome (1 or 0) and expectation (the presented probability of reward; Eq. 29). To rule out SPE as a driver of our results, we performed a confirmatory analysis, by constructing a GLM that included the model’s full aRPE signal and its SPE signal, and the compared the beta values of partial correlations associated with these regressors. For both the VTA/SN^73^ and the mPPC cluster, the average partial correlation weights for model’s full RPE was greater than for SPE signal (**Figure S11**).

### The hippocampus correlates with our computational model’s surprise that can enhance the value of anticipation

Our computational model predicts people receive enhanced anticipatory value following a surprise coincident with advance information cues. The magnitude of enhancement is proportional to the surprise, which is defined simply by the absolute value^59, 87^ of aRPE (Eq. 2 in the Method section). Our model predicts the boosting should be sustained over the entire duration of a wait period (**Figure 4a**), unlike the phasic (a)RPE signals that we just examined.^52, 88^

**Figure 4:**
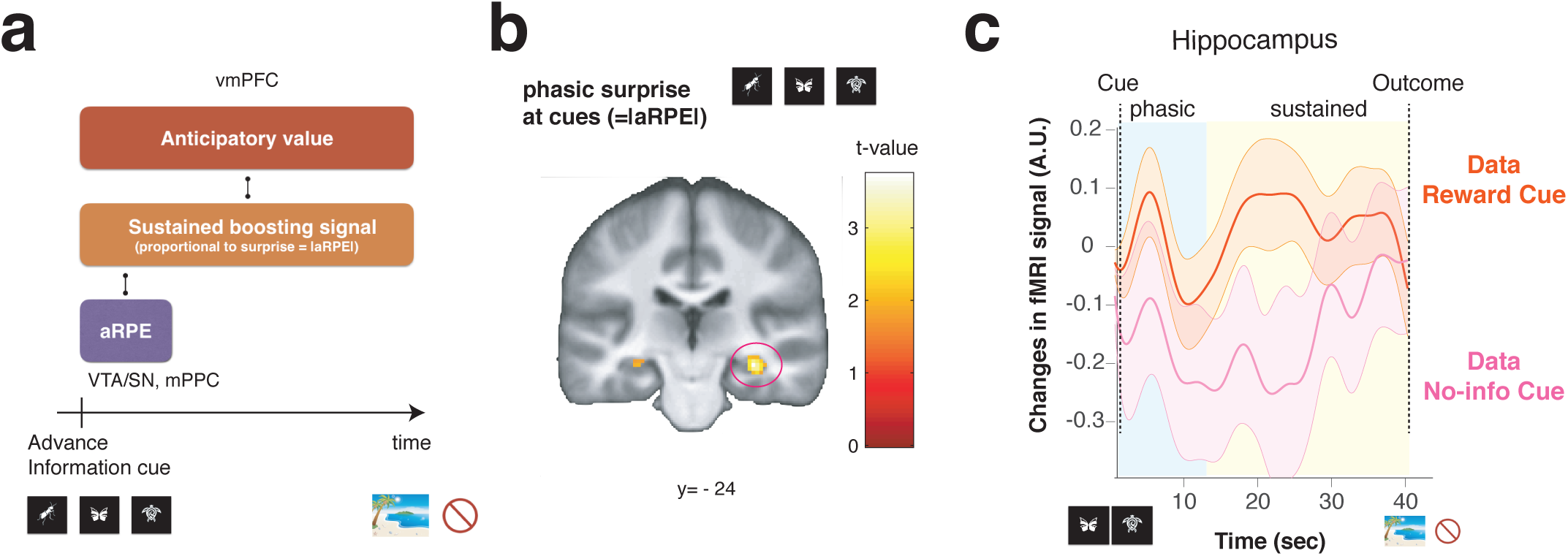
Neural correlates of our computational model’s surprise that can boost anticipatory values in our computational model. (**a**). Our model predicts that a surprise, quantified by the absolute value of aRPE, can boost the value of anticipation. The model predicts the effect of boosting to be sustained during the anticipatory period, in contrast to the phasic, short, aRPE signal, (**b**). A surprise at advance information cues, quantified by the absolute value of aRPE, significantly correlated with BOLD in the hippocampus (FWE *p* < 0.05 small volume correction^86^). (**c**). The temporal dynamics of fMRI signal in the hippocampus. Changes in activity averaged over participants after receiving a reward predictive cue (orange), and after receiving a no-information cue (magenta), are shown. The phasic response confirmed in (b) is apparent in the early phase of the delay period (blue), but the coding of boosting-related value are sustained over the course of entire wait period (blue and yellow), which is what our model predicted. The error bar indicates the SEM. Please also see **Figure S12** for responses to a no reward predictive cue.

Previous research suggests that the hippocampus is an ideal substrate for this effect. First, in the context of recognition tasks, the hippocampus encodes surprise (mismatch, novelty) signals.^36, 89–93^ Additionally, extensive studies implicate a couplings of hippocampus with the VTA/SN as well as with the PFC,^35–41, 47, 94^ the two regions that we we show are linked to our model’s computation. Further, although we do not specify psychological roots of our computational model’s enhancement of anticipatory value, we note that in the original study of anticipatory utility, the magnitude of anticipatory value is suggested to relate to the strength of imagination for future reward.^11^ Indeed, many studies link hippocampal activity to the imagination of future prospects (e.g.^42–44, 46^), where prefrontal-medial temporal interactions influence the effects of imagination on valuation,^45^ as well as support the mental construction of future events.^47, 48^

Therefore, we first examined phasic response of the hippocampus to a surprise at the onset of the advance information cue presentation, quantified by the absolute value of the model’s aRPE. As predicted, we found hippocampal activity was significantly correlated with the magnitude of a surprise (*p* < 0.05 FWE small volume correction by an anatomical mask of hippocampus;^86^ [32, −24, −12], *t* = 3.60, **Figure 4b**).

The phasic response to surprise is an important feature for the model’s boosting anticipatory value, but as outlined the model predicts that activity associated with boosting should be sustained until the ultimate reward delivery (**Figure 4a**). Therefore, we next asked whether the hippocampal response to a surprise, at advance information cues, persists throughout the delay period, as predicted by our computational model. Indeed, we found that hippocampal activity in the cluster that responded phasically to surprise at cue (the cluster is taken at *p* < 0.05 FWE small volume correction from the analysis in **Figure 4b**) was greater throughout the wait period after a reward predictive cue was presented (in which case a surprise was induced), compared to that following presentation of a no-information cue (in which case no surprise was induced), as seen in **Figure 4c** (see also **Figure S12** for responses to a no-reward predictive cue). Thus, in addition to expressing the magnitude of a surprise at advance information cues, hippocampal BOLD during the wait suggests features associated with our model’s signal that relates to boosting of anticipatory value.

### Midbrain-hippocampus-vmPFC circuit dynamically computes our computational model’s anticipatory value

So far we have shown that distinct regions encode our model’s computational signals. The vmPFC encodes our model’s value of anticipation; the VTA/SN (as well as the mPPC) encodes aRPE signal that is associated with a trigger for boosting of the value of anticipation, and the hippocampus encodes a sustained signal associated with our model’s boosting of the value of anticipation. In our computational model, these three signals are functionally coupled (please see **Figures 1ef, 4a** for schematic illustrations and Eqs 1 and 2 in the Method section for a more precise mathematical description). Specifically, as illustrated in **Figure 4a**, our model expects that a region that encodes a signal associated with sustained effect of boosting would be functionally coupled both to a region encoding aRPE and a region encoding the value of anticipation. Indeed, the hippocampal BOLD signal in **Figure 4c** suggests that it encodes both phasic (related to aRPE) and sustained (related to anticipatory value) signals. Furthermore, extensive studies implicate functional couplings of hippocampus with the VTA/SN as well as with the PFC.^35–41, 47, 94^

We hypothesized that sustained hippocampal activity mediates our model’s anticipatory value computation. In essence, in order to boost anticipatory values, the hippocampus links computations in the VTA/SN (aRPE) and the vmPFC (anticipatory value). To formally test this idea, we analyzed functional connectivity using dual psychophysical interaction (PPI) regressors based on two a priori seed regions: 1) the vmPFC (which encodes anticipatory value) and the model’s aRPE signal at advance information cues (which is encoded at the VTA/SN) as a psychological variable, and 2) the VTA/SN (which encodes aRPE) as a seed and the model’s anticipatory value signal (which is encoded at the vmPFC) as a psychological variable. Because each of these two PPI regressors includes variables relating to both the vmPFC (anticipation) and the VTA/SN (aRPE), and these variables are coupled in our computational model through the notion of boosting, this analysis tests our hypothesis that the hippocampus links the VTA/SN (aRPE) and the vmPFC (anticipation) as a potential medium of boosting. Thus, we included these two sets of regressors into the single GLM we used so far (see Methods section), and tested if hippocampal activity significantly correlated with these PPI regressors.

We found significant correlations in the hippocampus for both PPI regressors. Thus, the functional coupling between the VTA/SN (the area encoding aRPE) and the hippocampus^95^ was significantly modulated by our model’s anticipatory value signal (*p* < 0.05, FWE small volume correction;^86^ [22, −32, −6], *t* = 3.89, **Figure 5a**). Additionally, the functional coupling between the vmPFC and the hippocampus^96^ was significantly modulated by our model’s aRPE signal (*p* < 0.05, FWE small volume correction;^86^ [−30, −34, − 6], *t* = 3.70, **Figure 5b**.).

**Figure 5:**
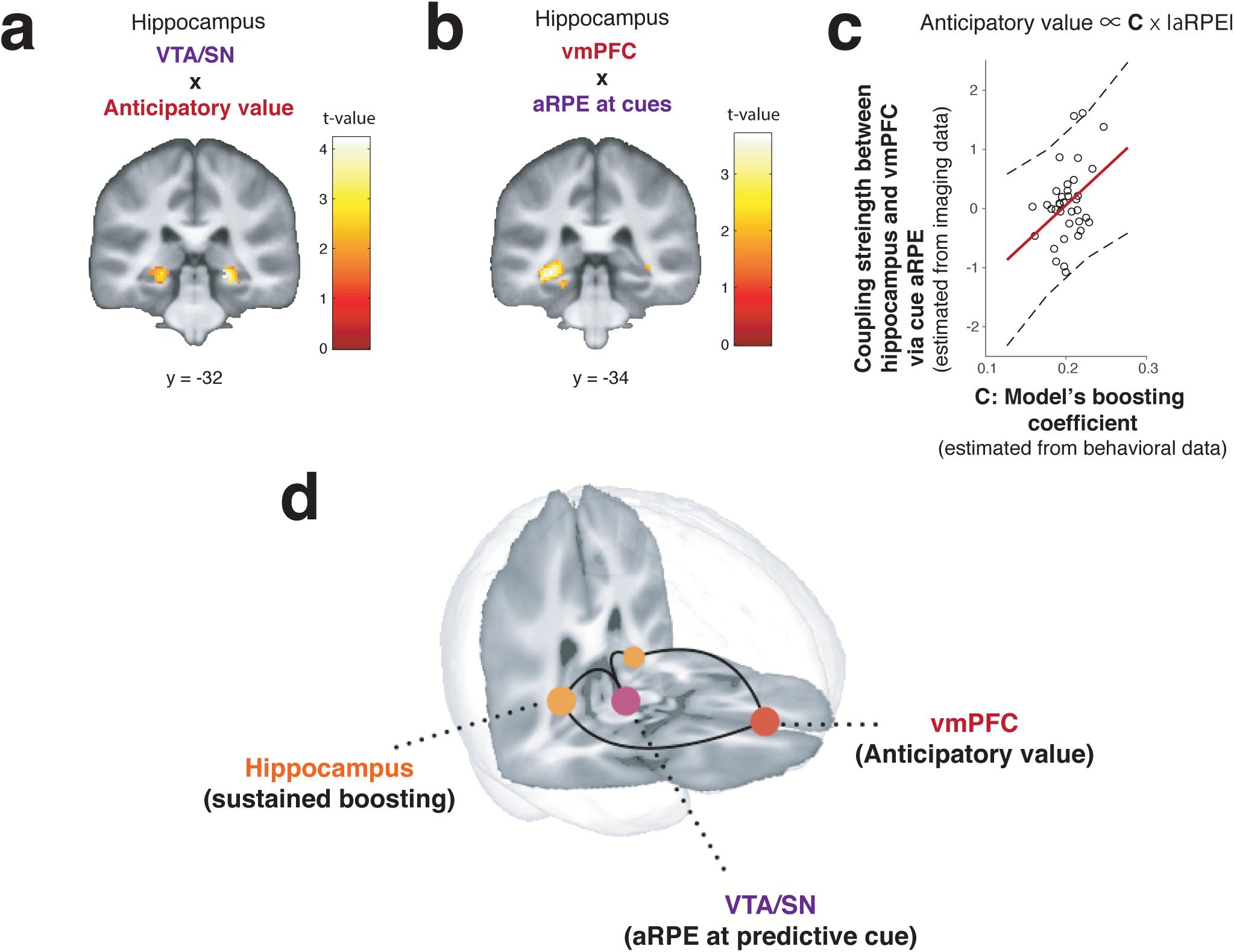
Functional connectivity analysis suggestive of a neural network associated with our model’s anticipatory value computation. (**a**). Functional coupling between the VTA/SN and the hippocampus is positively modulated by the model’s anticipatory value signal (*p* < 0.05 FWE small volume correction.^86^ PPI regressor: BOLD signal in VTA/SN modulated by model’s anticipatory value signal (encoded in vmPFC)). (**b**). Functional coupling between the vmPFC and the hippocampus is positively modulated by model’s prediction error signal (*p* < 0.05 FWE small volume correction^86^). PPI regressor: BOLD signal in vmPFC modulated by model’s prediction error signal. PPI regressors in (**a**) and (**b**) were chosen because each of them includes variables relating to both the VTA/SN (prediction error) and the vmPFC (anticipatory value), allowing us to test if the hippocampus links to both these two regions in a way predicted by our computational model. Note that (**a**) and (**b**) are partial correlation results from the same analysis using a single GLM with the two PPI regressors that are symmetrically orthogonalized. (**c**). The functional coupling strength between the vmPFC and the hippocampus mediated by the model’s prediction error signal is positively correlated with the model’s boosting coefficient parameter estimated by behavior of participants (*r* = 0.37, *p* < 0.05). This evidence suggests that our model’s predicted amplitude modulation of anticipatory value is mediated by the coupling between the hippocampus and the vmPFC, where the coupling strength is also modulated by our model’s aRPE encoded in the VTA/SN,^35^ as we predicted in our model. (**d**). Functional network for anticipatory value computation. Three distinctive regions play together to construct the anticipatory value, in a manner that is predicted by our computational model. The 3D brain image was constructed by the mean T1 brain images, which were cut at *y* = 34 and *z* = 15 for illustrative purposes.

If the hippocampal-vmPFC coupling is responsible for our computational model’s boosting of anticipation, the coupling strength that we estimated in our PPI analysis should relate to the model’s magnitude of boosting that we estimated from choice behavior. Our model predicts that the magnitude of boosting is linearly correlated with a parameter C, the linear boosting coefficient (Eq. 3, **Figure 5c**), which we had already fit to each individual participant. Therefore, we tested whether the linear boosting coefficient (that we estimated from our behavioral model-fitting) and the hippocampal-vmPFC coupling strength (that we estimated from our fMRI PPI analysis) are correlated with each other. As seen in **Figure 5c**, we found that these two variables estimated separately from imaging and behavioral data are positively correlated across participants. This is a strong evidence that links the three region network to our model’s anticipatory value computation. In particular, it suggests that the hippocampus-vmPFC connection, whose strength is influenced by our model’s aRPE that is encoded in the VTA/SN, is tied to our model’s enhanced strength of anticipatory value.

These functional connectivity results support our hypothesis that hippocampus plays a key co-ordinating role for our model’s computation. That is potentially boosting the value of anticipation, linking the vmPFC’s encoding of the value of anticipation with the VTA/SN’s encoding of prediction errors at advance information. The findings point to these regions functioning as a large-scale neural network for boosting the values of reward anticipation (**Figure 5d**), driving a preference for advance information in our task.

## Discussion

The value (utility) of anticipation has long been recognized as a key notion in behavioral economics and the cognitive sciences. While it has been linked to a wide range of human behavior that standard reward-value-based decision theories struggle to account for (e.g. a preference for advance information, risk-seeking, and addiction), the neural basis of the theory is unknown. Here, we took advantage of a new link between computational theory and behavior, and applied this perspective to fMRI data to reveal how the value of anticipation arises in the brain (please see **Figure S13** for a visual summary of this Discussion section).

Crucially, we show a network for computing the utility of anticipation, consisting of three specific brain regions. First, we show vmPFC represents the time course of an anticipatory value signal that evolved separately from a standard reward expectation signal during a waiting period. Second, dopaminergic midbrain regions, encompassing VTA/SN, encoded the model’s anticipation-reward prediction error (aRPE) that signals changes in expected value of anticipation and reward at advance information cues. Third, the hippocampus, whose activity indexed our model’s surprise signal, was functionally coupled both to the vmPFC and to the VTA/SN, in a sustained manner consistent with our model’s predicted boosting of anticipatory value. While the three-region’s functional coupling has been previously implicated in other settings,^35–41, 47, 94^ our study provides evidence for an explicit, mathematically defined, computational role. We suggest that its role in the contest of our study is to construct a value of anticipation which works as a reinforcement for behavior.

In turn, our study provides insights into neural processes underling human decision-making that standard decision theories struggle to explain. A case in point, in our current study, concerns a preference for early resolution of uncertainty,^14, 16, 18, 97, 98^ also known as information-seeking,^51–53, 99, 100^ or observing.^101, 102^ Humans and other animals are willing to incur costs to find out their true fate, even if this knowledge does not change actual outcome.^14, 53, 103–106^ An alternative idea, as opposed to our boosted anticipatory utility, is that people derive value from information itself.^52, 97, 107^ However this so-called intrinsic value of information cannot explain why a preference for advance information is valence-dependent,^53, 108^ that it depends on the reward probability in a way that does not covary with information-theoretic surprise,^109, 110^ and a sensitivity to delay until reward (as we also demonstrated here).^20, 54^ All of these findings are natural consequence of the utility of anticipation (but not of information per se).

Consequently, our results account for previous neural findings about the intrinsic value of information. This so-called information-prediction-error (IPE) signal is presumed to arise from the value of information,^51–53, 74, 75^ and has been reported in the same midbrain dopaminergic regions as standard reward prediction errors,^51, 52^ implying that the two signals might be strongly related.^111^ Indeed, our model accounts for information prediction errors as a side effect of anticipation-dependent aRPE, and we found that an aRPE signal positively correlates with BOLD signal in dopaminergic midbrain regions^51, 52^ as well as in the mPPC^81^ (see also^82^). We found that clusters in these regions are more strongly correlated with our model’s aRPE signal than with a standard RPE signal with no value of anticipation. This suggests that these regions encode our aRPE signal that unifies standard RPE signals as well as IPE signals.

Further, our results offer alternative accounts for addiction, and the possibility of individually-tailored psychiatric interventions (**Figure S13**). While initial phases of addiction^112^ involve excessive dopamine release at the time of drug consumption,^113^ later phases involve craving.^114, 115^ Our model can imply that people boost anticipatory value when the likelihood of drug administration increases (e.g. when purchasing drugs). People may feel greater value from obtaining drugs (which can act as a kind of conditioned stimuli^116^) than from administering them, because the former includes values associated with the anticipation of future administration. Importantly, our model predicts that people with certain parameter values (e.g. large boosting coefficients) could repeatedly over-boost the value of anticipating drugs, resulting in excessive, pathological, drug seeking (see Eq. 12). Although the learning process leading to pathological behavior may be very slow in a natural world, by fitting our model to subjects performing the task used here, we can in principle link an individual’s tendency toward addiction with unique cause of this disorder (e.g. excessive boosting or imbalance between anticipation and discounting). This can in turn suggest interventions tailored to individual patients, such as cognitive behavioral therapy focusing on controlling anxiety and craving,^117^ as well as possible dopaminergic antagonists to control boosting.

Our study also unifies separate notions concerning gambling: preference for risk and time-delay (the latter is called time-preference in behavioral economics). While these two economic phenomena have often been treated separately there is increasing evidence in favor of an interactive relationship (e.g.^118, 119^). Our computational model of anticipation explicitly offers an interaction between risk (prediction error) and delay (anticipation), because the former can enhance the value of latter. This interaction creates well-documented effects, such as nonlinear coding of probabilities of anticipated rewards.^120^ Thus our results suggests that neural computation in the three-region network can unify two separate economic preferences. It would be interesting to test our model’s predictions on how pharmacological manipulations (e.g. on dopamine) affect risk and time (delay) preference, where dopamine is likely to be heavily involved in computing aRPE. Further studies may allow us to design a behavioral task for psychiatric interventions, in which patients can lessen their preference for addicted substances, or even their risk preference in general, because our model can find the optimal task parameters for each individual to achieve this goal.

We found that hippocampus was involved in the value computation arising from reward anticipation, through its coupling with the VTA/SN and the vmPFC. Both hippocampus-VTA/SN and Hippocampus-(v)mPFC couplings have been extensively reported previously in animal studies as well as in some human studies (e.g.^35–41, 47, 94^). In rodents, hippocampus-PFC coupling has been shown to be gated by neurons in the VTA.^35, 39^ Oscillatory synchronization has also been reported in PFC-VTA-hippocampus axis in rodents performing a working memory task.^37^ Our finding is consistent with a previous finding in humans that activity of the VTA influences the baseline activity in posterior hippocampus.^94^ Posterior hippocampus, which we report in our PPI analysis, has also been linked to future simulation^121^ that is likely to relate to our model’s anticipatory value computation. Our functional connectivity analyses suggest that an aRPE signal encoded in the VTA/SN affects a functional coupling between the hippocampus and the vmPFC, which encode the enhancement of the value of anticipation; this can be tested in future studies involving pharmacological manipulations (e.g on dopamine). Because the hippocampus has a rich anatomical structure, further studies will illuminate how different parts of hippocampus contribute to value computation arising from reward anticipation.

Neuroeconomic studies show that people make decisions between goods in different categories, by expressing the value of those goods in a so-called common currency primarily encoded in the vmPFC (e.g.^122^). Here we found that the utility of anticipation expressed in the vmPFC. This invites an alternative interpretation of previously reported ramping activity in the vmPFC while waiting for rewards^123–125^ in terms of an anticipation-sensitive value signal, which has been interpreted as a reward-timing signal.

An alternative interpretation for our behavioral results is that participants do not like uncertainty. However, a previous study using the same task with aversive outcomes has shown that people avoid advance information when the outcomes are aversive.^100^ Another study also has shown a preference of advance information being valence dependent.^53^ These are consistent with our model’s predictions, but contradict simple uncertainty avoidance. In our model, advance information can boost negative anticipation for aversive outcome (i.e. dread^13, 55^), leading to an avoidance of (negative) advance information. Further studies will illuminate how advance information modulates dread in the brain, though hippocampal coding of sustained signal during a wait period for no-reward (**Figure S12**. Note: people assigned negative value to no-reward in our task, confirmed by our model fitting and self-reports) may suggest that a similar circuit presented here may be involved in the computation.

As is the case for the value of reward (whose psychological roots have been shown to be very complex^22, 61^), psychological roots of anticipatory value are likely to be complex. While we acknowledge that we had no control over what participants were thinking while waiting for outcomes in the scanner, participants’ informal self-reports after experiments were largely consistent with an idea that reward predictive cues made participants more excited while waiting for reward. We acknowledge that other psychological interpretations of our computational model are possible, as is the case for the roots of reward in a standard reinforcement learning model.^61^ For example, we should note an influential suggestion^16^ that future uncertainty drives other forms of anticipatory utility, such as anxiety. We did not consider this notion directly in our computational model; but in our model an agent can experience a mixture of positive and negative utilities of anticipation according to the probabilities of these outcomes (please see the Methods section). It would be interesting to study how this mixed anticipatory utility of our model relates to the notion of anxiety,^16^ which may help the design of more effective psychiatric interventions for anxiety disorders.

Finally, our study offers an alternative view to a long-standing problem in neuroscience and machine-learning. We refer here to the so-called temporal credit assignment problem which raises the issue of how neurons operating on a timescale of milliseconds learn relationships on a behaviorally relevant timescale (such as actions and rewards in our task). Designing a machine learning algorithm that overcomes this problem remains a challenge. Cognitively, our computational model suggests that the anticipation of future reward could serve as an aid to solve this problem, because a sustained anticipation signal can bridge the temporal gap between a reward predictive cue and an actual reward. A recent physiological study demonstrated that synaptic plasticity in hippocampal pyramidal neurons (e.g. place cells) can learn associations on a behaviorally relevant timescale, with the aid of ramping-like, slow, external inputs in a realistic setting.^126^ This has been shown to arise out of a slow input that can trigger a slow ramp-like depolarization of synaptic potential, which in turn unblock NMDA receptors, leading to a synaptic learning that spans duration of seconds.^126^ Our results thus suggest that a slow anticipatory value signal in the vmPFC that is sustained throughout long delay periods (or the sustained, coupled, activity in the hippocampus) could serve as such an input to neurons in the hippocampus, bridging the gap over behavioral timescales. A dopaminergic input from the VTA/SN to the hippocampus may facilitates this type of learning.^36, 127^ Further studies will illuminate how the learning takes place at a neuronal level.

In sum, we identified novel neural substrates for computing the value arising from anticipation, orchestrated by three distinctive brain regions with different associated functions. We suggest this anticipatory value drives a range of behaviors including information-seeking, addiction, and gambling. Our study can also provide a seed for individually tailored interventions for psychiatric disorders.

## Acknowledgment

We thank Tim Behrens for his insightful suggestion for the Fourier phase randomization test. We thank Colin Camerer, Okihide Hikosaka, George Loewenstein, Sandro Romani, Ethan Bromberg-Martin, Jackie Gotlieb, Elliot Ludvig, Yunzhe Liu, Bastian Blain, Giles Story, Laurence Hunt, Jeff Cockburn, Vincent Man, Tomas Aquino, Caroline Charpentier, Bowen Fung, Wolfgang Pauli, Erin Burkett for the most valuable discussions and helpful suggestions for the manuscript. We also thank radiographers at the UCL for their assistance in running fMRI experiments. This work was supported by the Max Planck Society, the Gatsby Foundation, Wellcome Trust Investigator Award, Japan Society for the Promotion of Science, the Swartz Foundation, Wellcome Sir Henry Dale Fellowship (211155/Z/18/Z), the Jacobs Foundation (2017-1261-04), the Medical Research Foundation, and 2018 NARSAD Young Investigator grant (27023) from the Brain & Behavior Research Foundation.

## Methods

### Participants

Thirty-nine self-declared heterosexual male participants were recruited from the University College London (UCL) community. Participants provided informed consent for their participation in the study, which was approved by the UCL ethics committee.

### Experimental task

The task was a variant of that in,^20^ which itself was inspired by a series of animal experiments into information-seeking or observing behavior (e.g.^51, 54^). At the beginning of each trial, a pair of task-information stimuli (hourglass and partially-covered human silhouette) were shown along with two choice targets. The number on the hour-glass indicated how long the participants had to wait until seeing a reward or no-reward, where 1/2, 1, 2 4, 8 hour-glass meant 1, 5, 10, 20, 40 sec of waiting time, respectively. The other stimulus, a partially-covered human silhouette, indicated the probability of seeing a reward, specified by the area of uncovered semi-circle (5, 25, 50, 75, 95 % chance of rewards). Two lateral rectangular targets were presented as choices: the Immediate-Information target marked as ‘find out now’, and the no-information target marked as ‘keep it secret’. The positions of the hourglass, and the covered silhouette, were kept the same every trial, but the locations of choice targets were randomly alternated between left and right on each trial.

The participants were required to choose between left and right targets by pressing a button within three seconds. Once the participants chose a target, one of the three cues appeared in the center of the screen. If the participants chose the Immediate-information target, then a cue that signalled upcoming reward or no-reward appeared on the screen until the onset of reward or no-reward. If the participant chose the No-information target, a cue that signalled no-information about reward appeared on the screen. The meaning of the cues were fully instructed to participants beforehand. The meanings of the cues were counter-balanced across subjects. In order to ensure immediate consumption, rewards were images of attractive female models from a set that had previously been validated as being suitably appetitive to heterosexual male subjects;^20, 50^ reward images were presented for 1s. Images were chosen randomly from the top 100 highest rated pictures that were introduced in.^50^ No image was presented more than twice to the same participants. In case of no-reward, an image signaling absence of a reward was presented for 1s. In either case, a blank screen was presented for 1s before starting a new trial. These timings were set to reduce the timing uncertainty which may cause prediction error that can interfere with our model’s value computation.^83^

Participants were fully instructed about the task structure including the meaning of stimuli about the probability and delay conditions, as well as the advance information cues. Then participants underwent extensive training that consisted of three tasks: a variable-delay but fixed probability task, a fixed-delay but variable probability task, a variable-delay and variable-probability task. This ensured that participants had fully learned the task and had adequately developed preferences before being scanned. Scanning was split into three separate runs, each of which consisted of 25 trials that covered all conditions once. Trial orders were randomized across participants. Subjects had a break of approximately 30 sec between runs.

### Computational model

The model for the task is fully described in.^20^ Briefly, following Loewenstein’s suggestion that the anticipation of rewards itself has hedonic value^11, 13^ (e.g. subjects enjoy thinking about rewards while waiting for them), we extended a standard reinforcement learning framework to include explicit reward anticipation, which is often referred to as savoring.^11^ The model’s innovation is to suggest that the value of anticipation can be boosted by reward prediction errors associated with advance information about upcoming rewards.^20^ We note that savoring here is a mathematically defined economics term, and is different from (though may be related to) savoring in positive psychology (the acts of enhancing positive emotions).^128^

To describe the model formally, consider a task in which if a subject chooses the Immediate-Information target, then they receive at *t* = 0 a reward predictive cue *S*^+^ with a probability of *q*, or a no-reward predictive cue *S*^−^ with a probability of 1 −*q*. Subsequently, the subject receives a reward or no-reward at *t* = *T* (= *T*_Delay_), with a value of *R*^+^ or *R*^−^, respectively. In our recent experiment we found that subjects assigned a negative value to an absence of reward;^20^ but this is not necessary to account for preference for advance information that has been observed in animals.^15, 54^

Based on the observation that subjects prefer to delay consumption of certain types of rewards, Loewenstein proposed that subjects enjoy anticipation while waiting to enjoy the actual reward.^11, 13, 55^ Formally, the anticipation of a future reward *R*^+^ at time *t* is worth 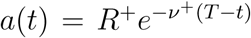, where *ν*^+^ governs its rate. Including *R* itself, and taking temporal discounting into account, the total value of the reward predictive cue, 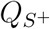, is

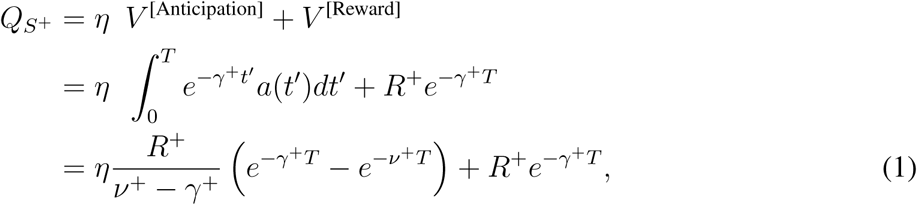

where *η* is the relative weight of anticipation, *γ*^+^ is the discounting rate, and *T* is the duration of delay until the reward is delivered. In prior work, *η* had been treated as a constant that relates to subjects’ ability to imagine future outcomes^56^; however, we proposed that it can vary with the anticipation-reward prediction error *δ*_aRPE_ (aRPE) at the time of the predicting cue.^20^ Our proposal was inspired by findings of the dramatically enhanced excitement that follows such cues.^54^ A simple form of boosting arises from the relationship

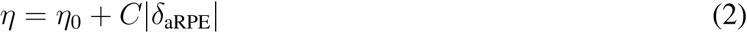

where *η*_0_ specifies the base anticipation, and *C* determines the gain. That anticipation is boosted by the *absolute value* of aRPE is important in applying our model to comparatively unpleasant outcomes.^20^ We also note that the boosting is expected to be sustained throughout a wait period, unlike phasic RPE (or aRPE) signals.^23^

The total value of the no-reward predictive cue, 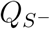, is then

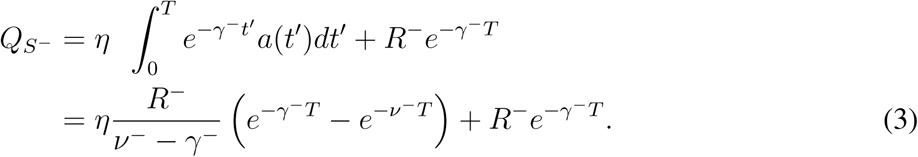

Following our previous work, we assumed that *γ* = *γ*^+^ = *γ*^−^.

Note that in our model, aRPE affects the total cue values 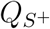 and 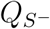, which in turn also affect subsequent aRPEs. Therefore the linear ansatz for the boosting of anticipation by aRPE (Eq.(2) could lead to instability due to unbounded boosting. This instability could account for maladaptive behavior such as addiction and gambling. However, in a wide range of parameters, this ansatz has a stable, self-consistent, solution. In our experiment, the prediction error for the reward and no-reward predictive cues can be expressed as.

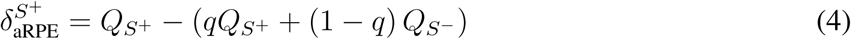

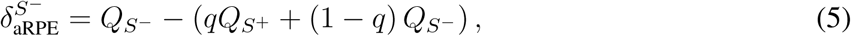

which are, assuming the linear ansatz,

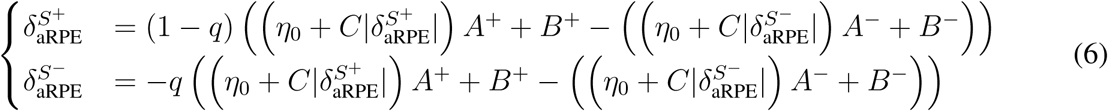

where

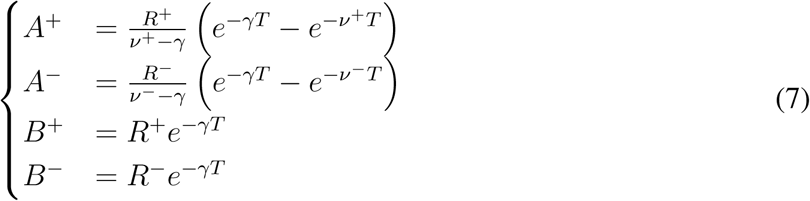

Assuming that *R*^−^ ≤ 0 and 0 ≤ *R*^+^, Equations (6) imply that 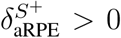 and 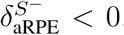. With this, Equations (6) can be reduced to

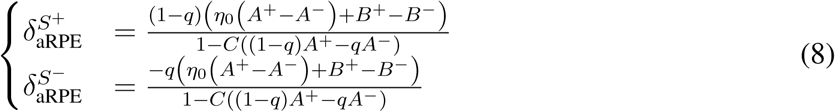

Because (*η*_0_ (*A*^+^ *− A*^−^) + *B*^+^ *− B*^−^) > 0, in order that 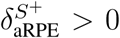 and 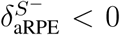 hold for all *q* and *T*, the denominators must be positive for all 0 ≤ *q* ≤ 1 and 0 ≤ *T*. In other words,

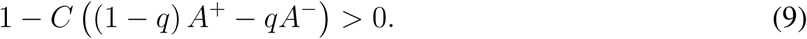

for 0 ≤ *q* ≤ 1 and 0 ≤ *T*, meaning that

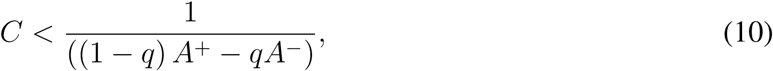

for 0 ≤ *q* ≤ 1 and 0 ≤ *T*. This means that

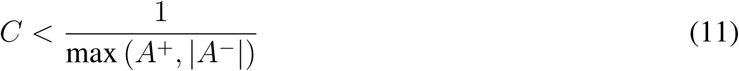

for 0 ≤ *T*. It is straightforward to show that *A*^+^ takes its maximum at 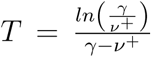, and |*A*^−^| at 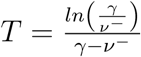. Thus the condition that the linear ansatz gives a stable self-consistent solution is

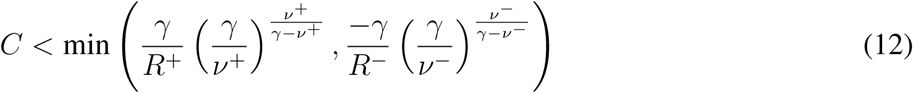

In our model-fitting, we imposed this stability condition. Violating it could account for maladaptive behavior such as addiction and pathological risk seeking.

An alternative to imposing such a stability condition would be to assume that boosting saturates in a non-linear manner:^20^

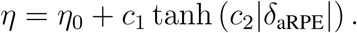

>However, the model’s qualitative behavior does not depend strongly on the details of the aRPE dependence of anticipation.^20^ Hence we only used the linear ansatz in our analysis in the current study.

We note that our current computational model^20^ was originally introduced in light of an experimental refutation^52^ of an earlier reinforcement-learning model for information-seeking (observing) behavior.^102^ This early computational model^102^ appealed to the same underlying idea discussed here, namely that of a Pavlovian influence^129^ of a prediction error signal over actions.^130^ The present model considers an anticipatory process (a type of an internal action^130^), in contrast to this earlier model that characterized this effect in terms of suppressed forgetting.

We generated choice probability^131^ from our model by taking a difference between the expected value of immediate information target and that of no-information target and taking it through sigmoid with a noise parameter *σ*.^20^

For our model comparison, we also fit a model with no anticipation *η* = 0, and a model with an anticipation but that is not boosted by aRPE, i.e. *C* = 0.

### Behavioral Model fitting

We used a hierarchical Bayesian, random effects analysis.^20, 49, 132^ In this, the (suitably transformed) parameters **h**_*i*_ of participant *i* are treated as a random sample from a Gaussian distribution with means and variance ***θ*** = {***µ***_*θ*_, **Σ**_*θ*_} characterising the whole population of subjects; and we find the maximum likelihood values of ***θ***.

The prior distribution ***θ*** can be set as the maximum likelihood estimate:

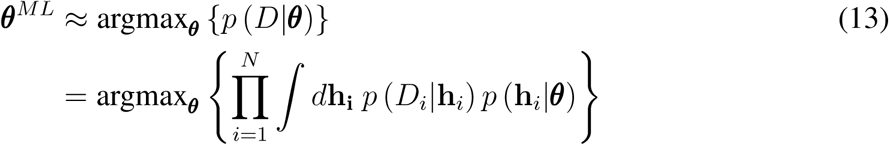

We optimized ***θ*** using an approximate Expectation-Maximization procedure. For the E-step of the k-th iteration, a Laplace approximation gives us

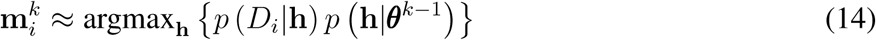

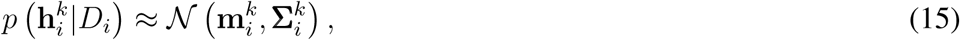

where 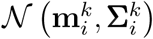 is a Normal distribution with mean 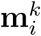 and covariance 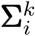 that is obtained from the inverse Hessian around 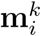. For the M step:

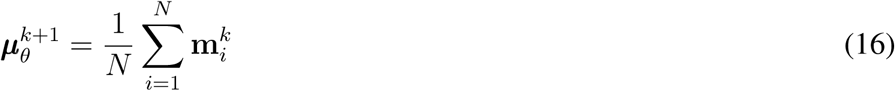

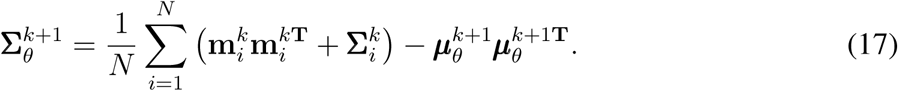

For simplicity, we assumed that the covariance 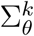 had zero off-diagonal terms, assuming that the effects were independent.^49^

### Model comparison

We compared the goodness of fit for different computational models according to their integrated Bayes Information Criterion (iBIC) scores.^20, 49^ We analyzed log-likelihood of data *D* given a model *M*, log *p*(*D*|*M*):

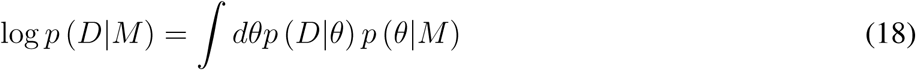

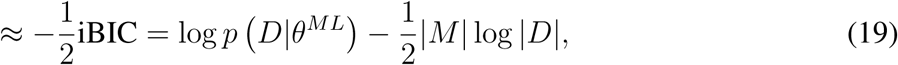

where iBIC is the *integrated* Bayesian Information Criterion, |*M*| is the number of fitted prior parameters and |*D*| is the number of data points (total number of choice made by all subjects). Here, log *p*(*D*|*θ*^*ML*^) can be computed by integrating out individual parameters:

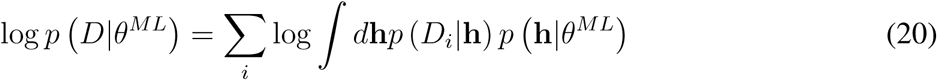

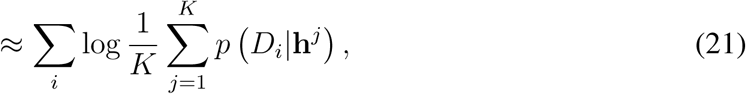

where we approximated the integral as the average over *K* samples **h**^*j*^’s generated from the prior *p* (**h**|*θ*^*ML*^).

### Model’s fMRI predictions

Our computational model makes specific predictions about temporal dynamics of anticipatory, reward, value signals during wait periods, as well as unique anticipatory reward prediction error (aRPE) signals at predictive cue onsets. Using the parameters (MAP estimates) for each participant, we generated the following variables for each participant as parametric regressors for the fMRI analysis.

The temporal dynamics of anticipatory value signal for positive domain at time *t* during wait period, until reward onsets *t* = *T* are:

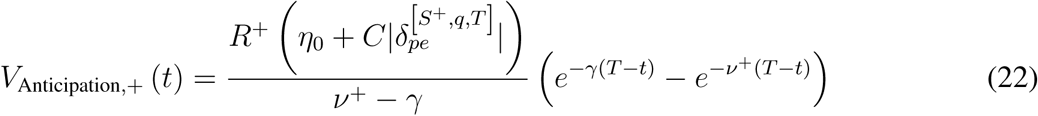

for the negative domain:

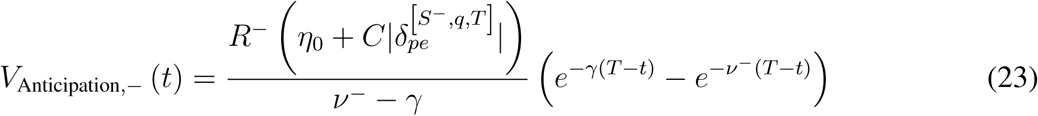

We expressed these as two separate regressors.

Because aRPEs explicitly enter the value function of the immediate information via boosting, aRPE and the value of the immediate information target influence with each other, needed to be computed in a self-consistent manner (Eq. 5). We assumed the consistency was achieved for participants through their extensive training sessions. The aRPE 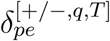 are determined for each delay *T* and reward probability *q* condition self-consistently (see below). After a no-information choice, these signals are scaled by the probability of reward *q* or no-reward 1 −*q* (and no prediction errors). Note that we set *R*^+^ = 1 without loss of generality.

The discounted reward signal at *t* during the wait period is expressed as

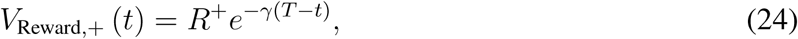

while the discounted no-reward signal at *t* is

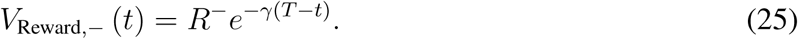

Note that the anticipatory value signal is an integral of (discounted) anticipation urgency signal:

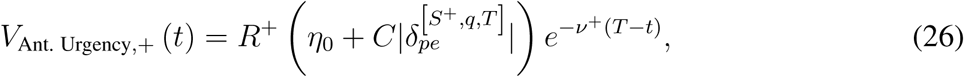

and

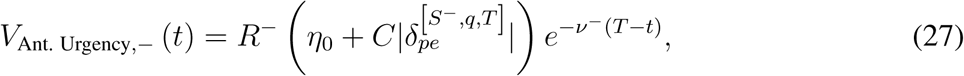

which we also included to the GLM.

The aRPE at information cue onsets are computed for each condition (*q, T*) self-consistently according to Eqs.(8). That is

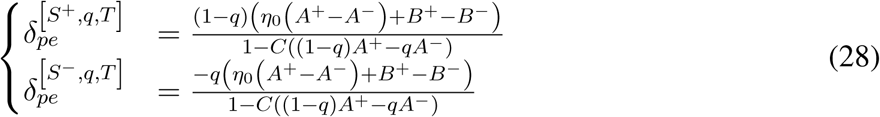

where *A*^+*/−*^, *B*^+*/−*^, are given by Eqs.(7). In our analysis we put positive and negative aRPE as a single parametric regressor at information cue onsets. Because the aRPE is expressed as the difference between the model’s presented cue value and the model’s expected cue value Eq.(5), we also tested a region is positively correlated with the model’s presented cue value and negatively correlated with the model’s expected cue value in Eq.(5).

Note that the aRPE signal is different from other conventional prediction error signals, including the so-called state prediction errors:^85^

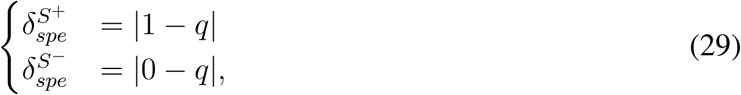

and a standard RPE signal with reward value alone (we can obtain this by setting *C* = *η*_0_ = 0 in Eq.(28)):

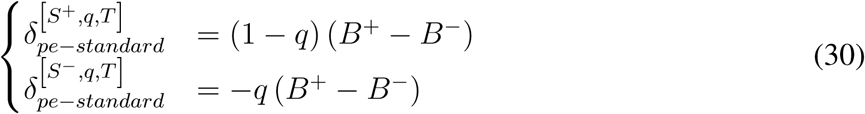

which we used for a control analysis.

### fMRI data acquisition

We acquired MRI data using a Siemens Trio 3Tesla scanner with a 32-channel head coil. The EPI sequence was optimized for minimal signal dropout in striatal, medial prefrontal, and brainstem regions:^133^ 40 slices with 3-mm isotropic voxels with repetition time (TR) 2.8 s and echo time (TE) 30 ms, and slice tilt 30 degrees. Additionally, field maps (3-mm isotropic, whole-brain) were acquired to correct the EPIs for field-strength inhomogeneity.

### fMRI analysis

We used SPM12 (Wellcome Trust Centre for Neuroimaging, UCL, London) for standard preprocessing and image analysis. The first 5 scans of each session were discarded to account for T1-saturation effects. The standard preprocessing includes: slice-timing correction; realigned and unwarped with the field maps that were obtained before the task; co-registration of structural T1 weighted images to the sixth functional image of each subject; segmenting structural images into grey matter, white matter and cerebral spinal fluid; normalising structural and functional images spatially to the Montreal Neurological Institute (MNI) space; spatially smoothing with a Gaussian kernel with full-width half-maximum of 8mm. The motion correction parameters were estimated from the realignment procedure, and were included to the first level GLM analysis.

We regressed fMRI time series with GLMs that consist of onset regressors and our model’s signals, as well as nuisance regressors. Onset regressors included the presentations of the initial screen, the presentations of cues, the presentation of outcomes. The parametric modulated regressors were added to the presentation of cues with model’s aRPE, and outcome onset with the value of reward or no reward. The onsets of cues preceding the shortest delay (1*s*) was separately modeled so that the prediction errors at the cues were not affected by reward. The time course of our model’s signals included the anticipatory value signals for positive and negative domains, the discounted value signals for positive and negative domains, anticipatory urgency signal for positive and negative domains. The model’s predictive signals were generated for each of the anticipatory periods, using the model that was fit to each participant, which were then convolved with the canonical HRF function. We added nuisance parameters that consist of movement estimated from preprocessing, large derivatives of movement between volumes that were larger than 1 mm, box function during the anticipatory periods, box function for each experimental run. In our control analysis, we also added box function during the anticipatory periods that was parametrically modulated by constant expectation of reward, parametrically modulated cue presentation with state prediction errors.

### Regions of interests

The region of interests for VTA/SN was taken from.^73^ The region of interests for hippocampus was taken from.^86^

### PPI analysis

We performed PPI analysis with a single GLM, which contained 1) BOLD signal of VTA/SN 2) a PPI regressor that is an interaction between the BOLD signal of VTA/SN and model’s anticipatory value signal 3) the BOLD signal of vmPFC 4) a PPI regressor that is an interaction between the BOLD signal of vmPFC and model’s aRPE signal, as well as other onset/movement regressors that we included in our original analysis.

The vmPFC’s seed activity was defined as the first eigen value of the BOLD signal in the cluster correlated with our model’s anticipatory value (*p* < 0.05 whole brain FWE corrected), and VTA/SN’s seed activity was defined as the eigen value of the BOLD signal in the cluster that was significantly correlated with our model’s aRPE at cues (*p* < 0.05 FWE small volume corrected^73^). Note that both aRPE and anticipatory value signals were controlled in our PPI analysis.

## Availability

The data and codes that support the findings of this study are available from the corresponding author upon reasonable request.

## Supplemantary figures

**Figure S1:**
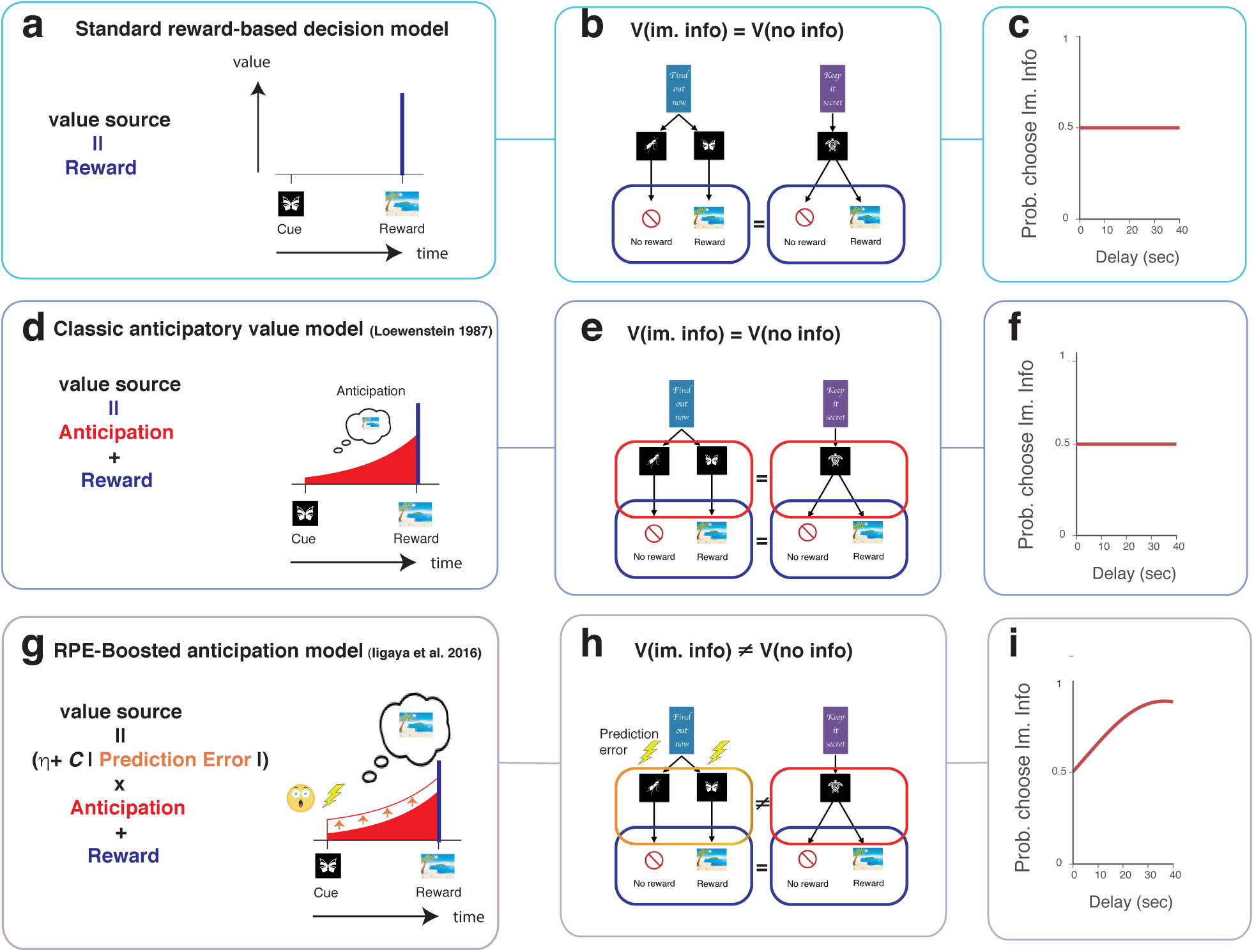
The mechanism by which the model predicts increasing preference for advance information, as a function of the duration of the wait period. (**a**,**b**,**c**). A classical reward-based decision-making model. This class of models assumes that a choice is made according to the attractive value of discounted future reward (a). Because there is no difference in the probability of obtaining reward between the Immediate-info-target and the No-Info-target, the model assigns the same value to the two choice targets (b). As a result, the model predicts no preference between the two targets across different delay conditions (c). (**d**,**e**,**f**). The classical behavioral economic model of the utility of anticipation.^11^ The model assumes that people experience value from the anticipation of future reward, in addition to from consumption of the reward itself. Although this model can capture the well-documented behavior that subjects delay reward consumption, it assigns the same values to the Immediate-info-target and No-info-target in our task (e). As a result, this model also predicts indifferent choice between the two targets (f). (**g**,**h**,**i**). aRPE-boosting anticipation model.^20^ Inspired by an observation of a dramatic increase in excitement after receiving the information that resolves uncertainty about upcoming reward,^54^ this model hypothesizes that the value of anticipation can be boosted by prediction errors associated with the reward predictive cues (g). Consequently, the value of the Immediate-info target can become greater than the value of the No-info target, as the duration of the wait becomes longer (h). As a result, the model predicts that subjects show stronger preference of the Immediate-info-target in longer delay conditions. This model has been previously validated in a series of behavioral experiments^20^ (i).

**Figure S2:**
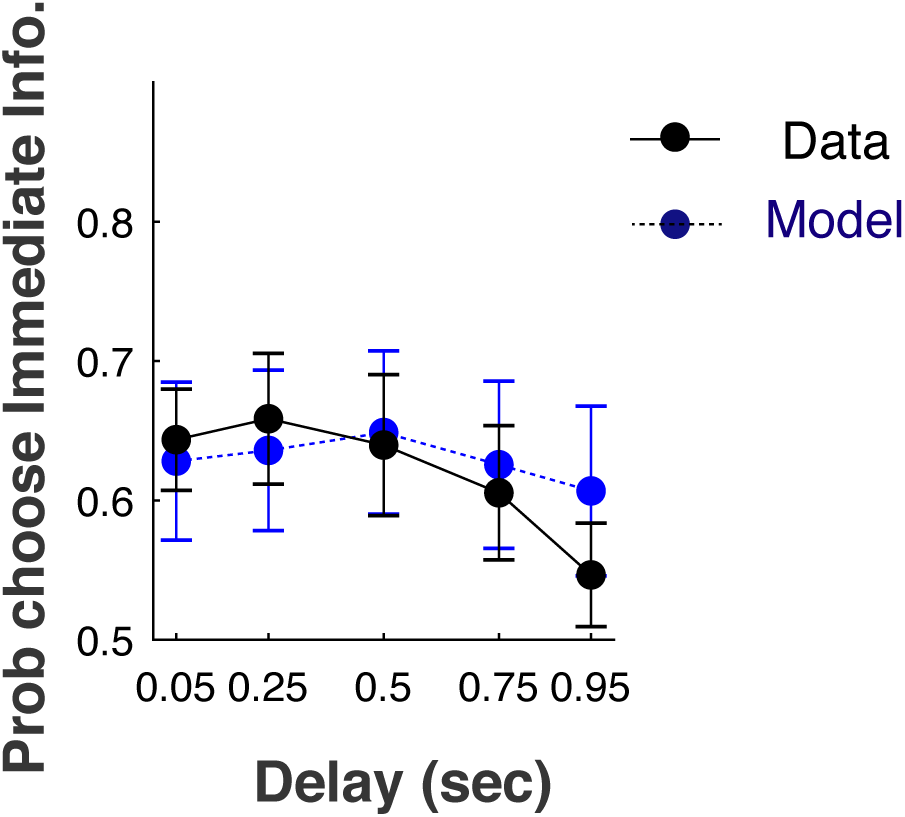
Our model (blue) captured choice preference over reward probability conditions in data (black). The error bars indicate the SEM of participants (n=39).

**Figure S3:**
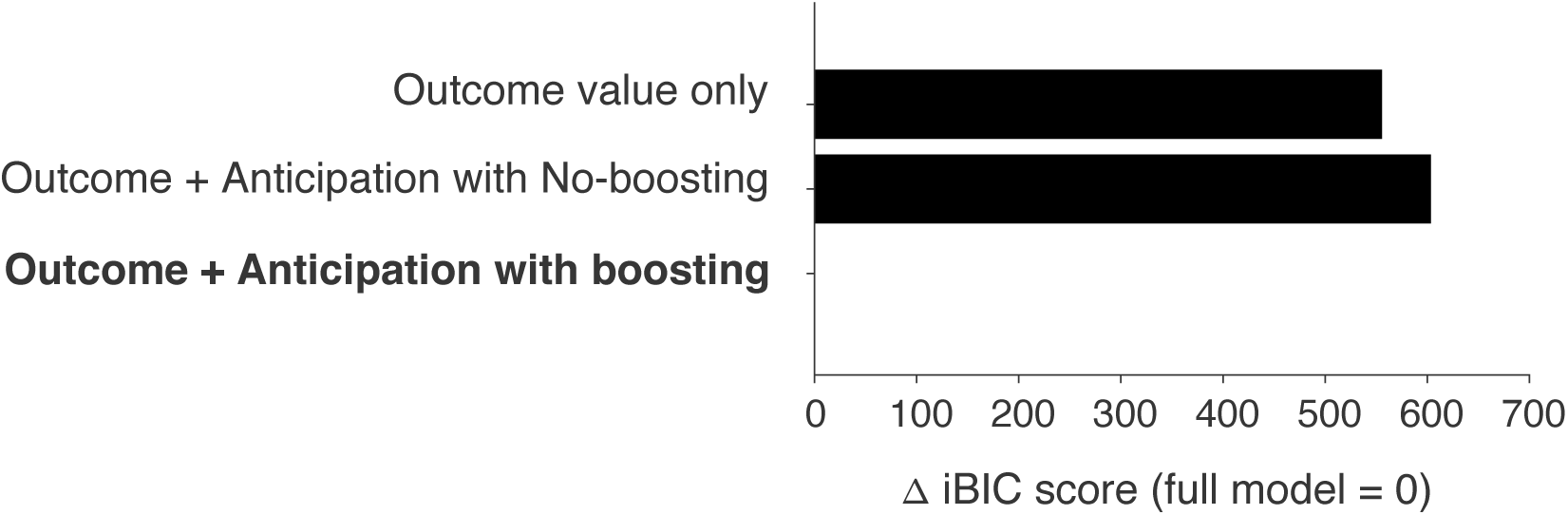
Our model comparison using integrated Bayesian Information Criterion (iBIC) strongly favors our full model with values of outcomes and aRPE-boosted anticipation, over a model with outcome values but no anticipatory utility, as well as a model with values of outcomes and anticipation that is not boosted by aRPE. All models included temporal discounting. A smaller score indicates a better model. The score is shown in log-scale. Please see the Methods sections for the precise definitions of the models.

**Figure S4:**
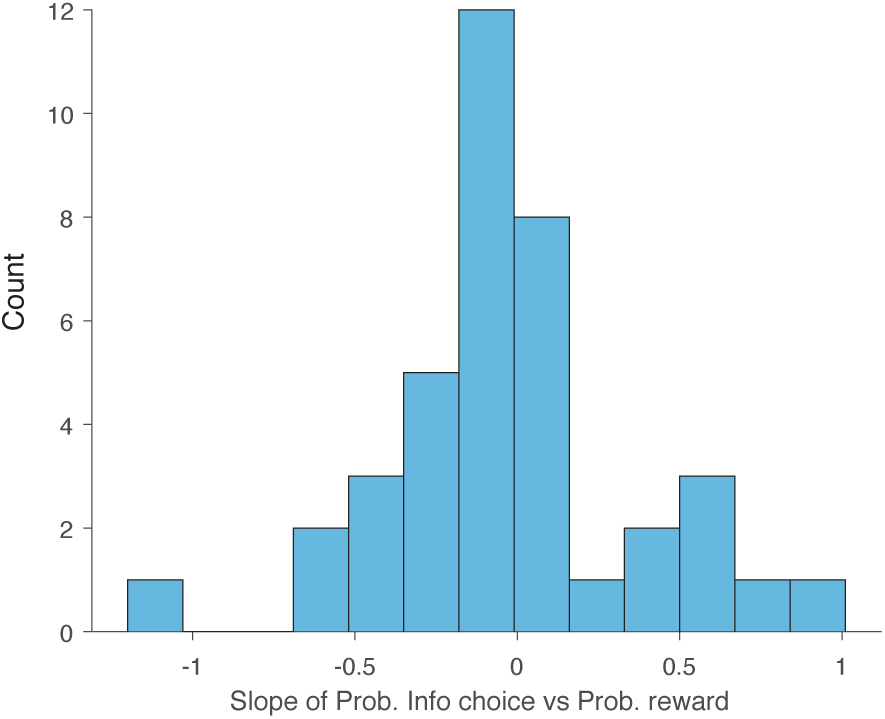
Heterogeneity amongst the participants in how their preferences for Info-target depended on the reward probability. The plot shows a histogram across the subjects of the slope of a linear fit to the probability of choosing the Info-target versus the probability of reward.

**Figure S5:**
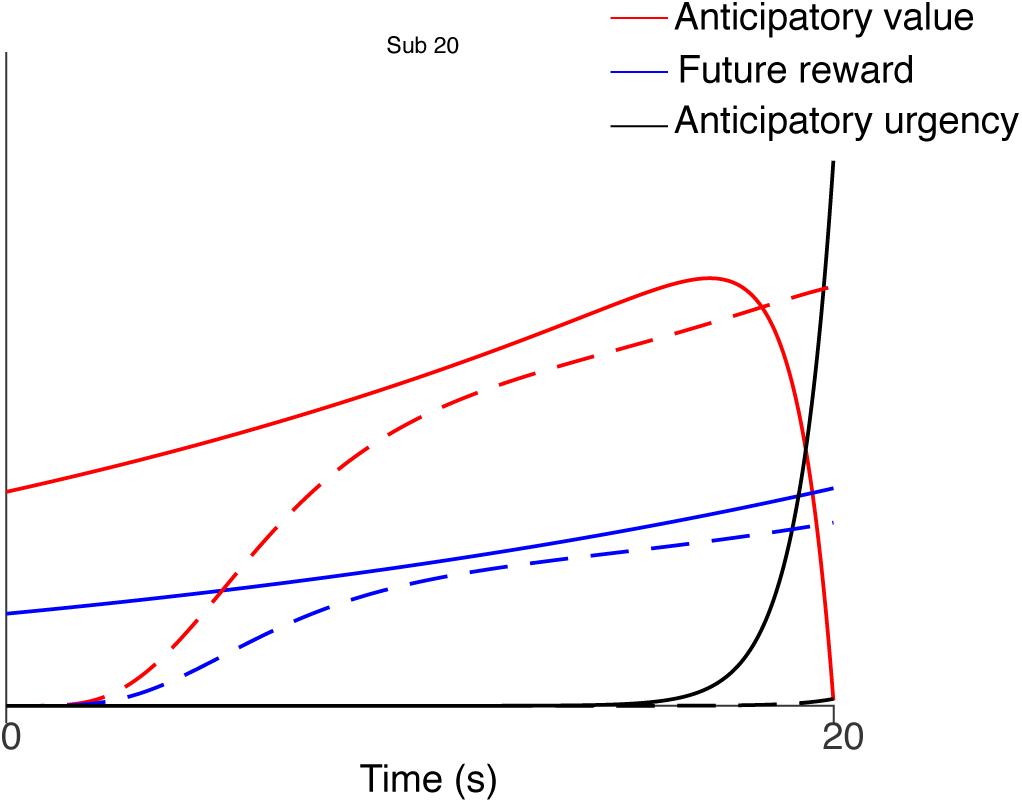
Prediction of fMRI signal in one subject (subjust 20). The anticipatory value signal (red). The discounted reward signal (blue). The anticipation stream signal (black). The dashed curves indicate HRF convoluted predictions for fMRI.

**Figure S6:**
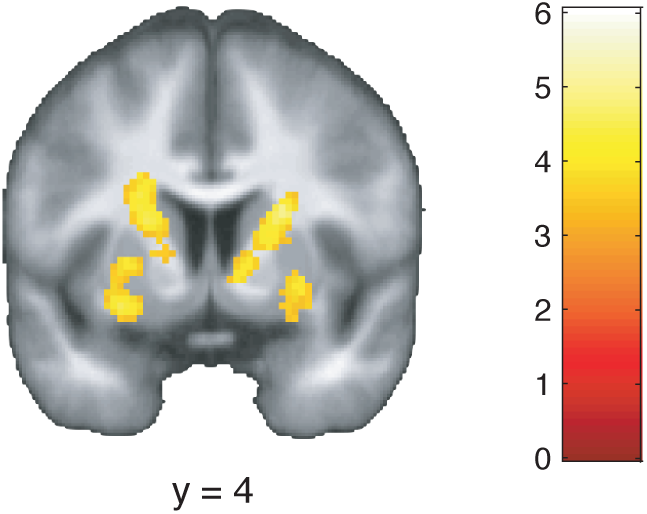
Voxels in dosal caudate correlated with anticipatory value signal. The effects in caudate survived our phase-randomization test (*p* < 0.001 whole-bran FWE correction). The effects in posterior putamen did not survived the whole-brain correction. Voxels at *p* < 0.001 (uncorrelated) are shown for display purposes.

**Figure S7:**
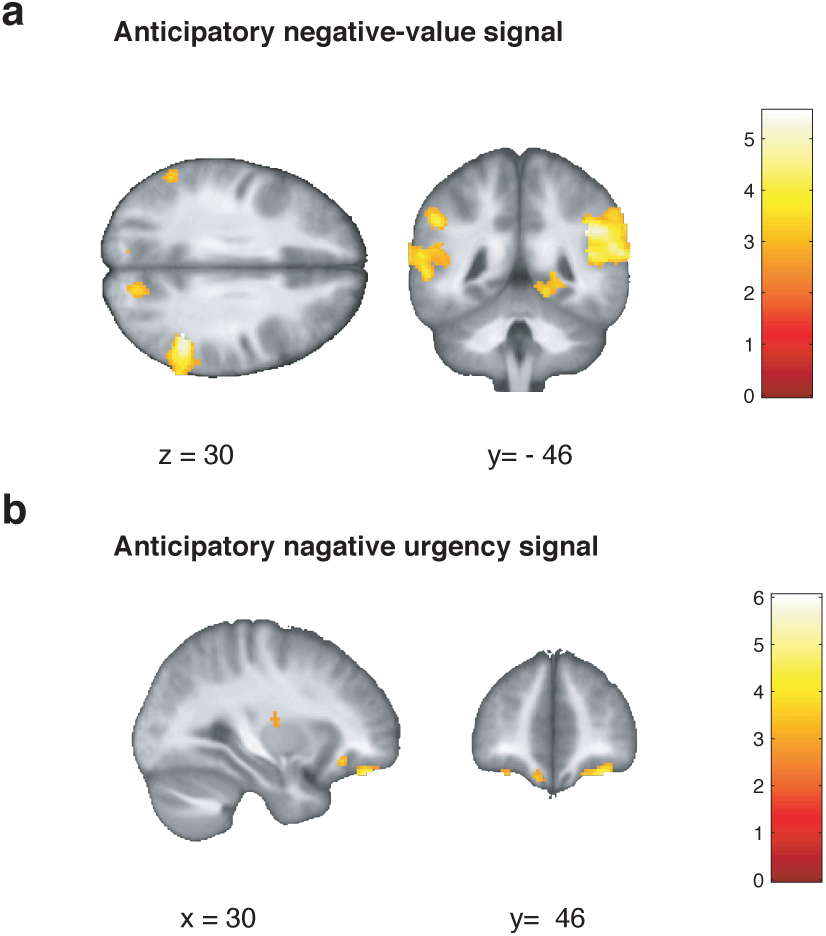
fMRI correlates in negative (no-reward) domain. (**a**) Correlation with anticipatory value signal for no-reward, which is in the negative domain. (**b**) Correlation with anticipatory urgency signal in the negative domain. In all panels, voxels at *p* < 0.005 uncorrected are shown for display purposes.

**Figure S8:**
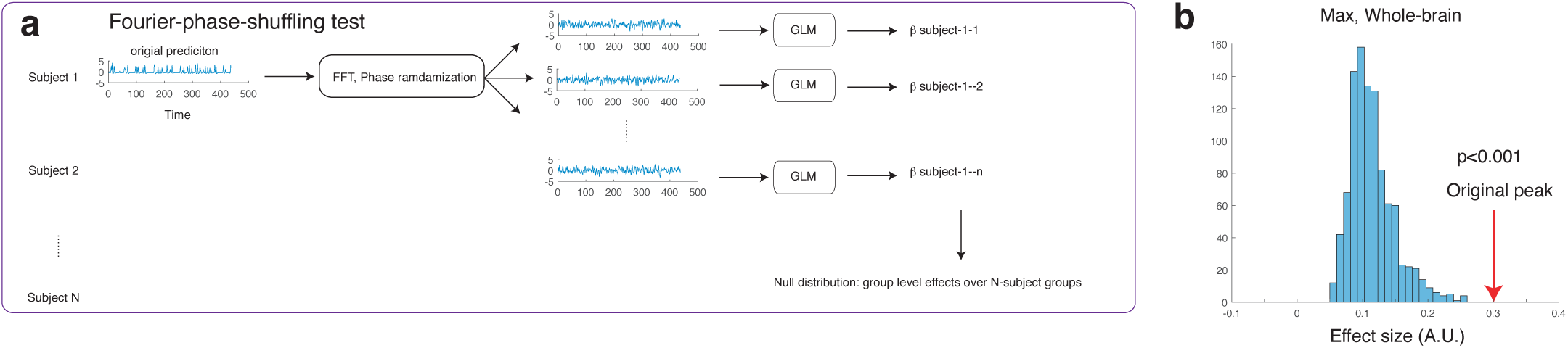
Our control analysis using phase-randomization validates the correlation between the model’s anticipatory value signal and the fMRI signal. (**a**) Schematics of the analysis. In response to recent article about false-positive correlations between slow signals in neuroscience,^69^ we performed a new analysis using phase-randomization of signals. For this, we first transformed our model’s predicted anticipatory signal into the Fourier space. Then we randomized the phase of each frequency without disturbing the power, before transforming back to the original space. We then ran the standard GLM analysis using this regressor as a model’s prediction to estimate the regression coefficient. We repeated this for each participant over 100 times (3,900 GLMs in total). We then randomly selected GLM results over participants (one from each participant) to perform a standard second level analysis. We repeated this second level analysis for 1,000 times to create a null distribution of the effect. The null distribution was constructed by taking the maximum correlation over each GLM result, and this was compared against the original. (**b**) Our test shows that our original correlation is significantly greater than by chance, compared to the null distribution constructed by the phase-randomization method (*p* < 0.001)

**Figure S9:**
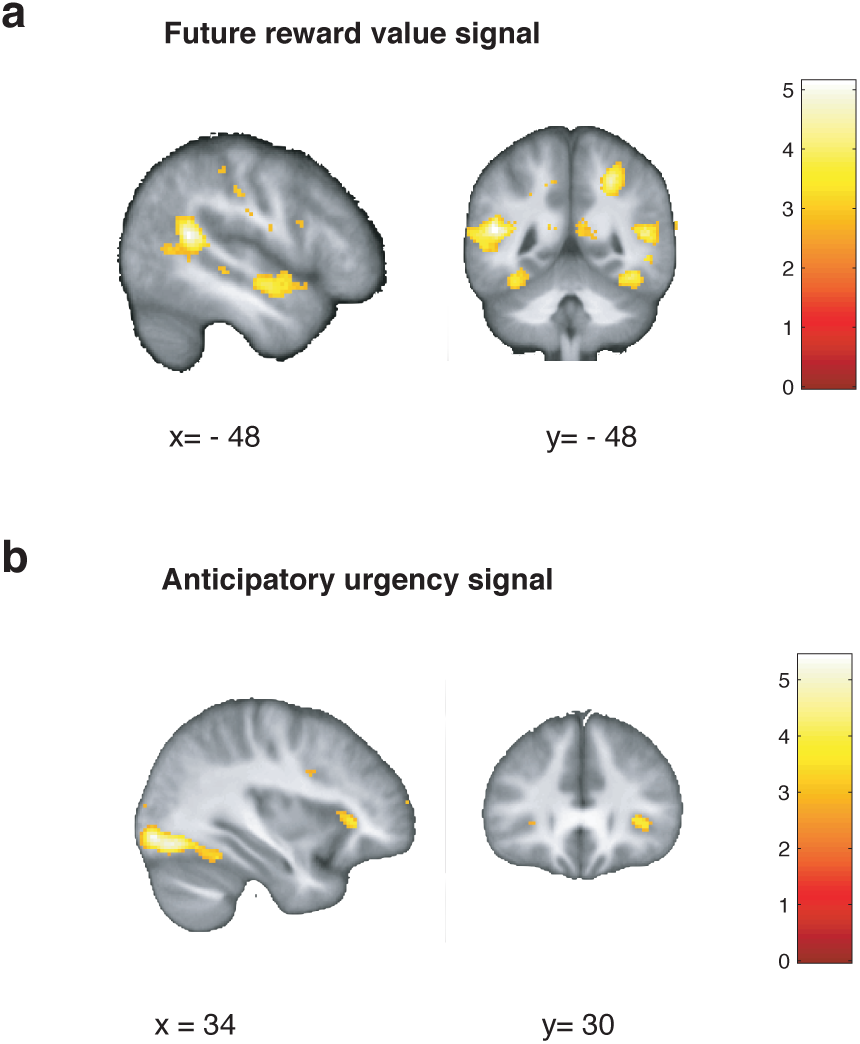
(**a**) Correlations with discounted future reward signal. Regions in superior temporal gyrus survives the whole brain FWE correction *p* < 0.05. (**b**) Correlations with anticipatory urgency signal. In all panels, voxels at *p* > 0.005 are shown for display purposes.

**Figure S10:**
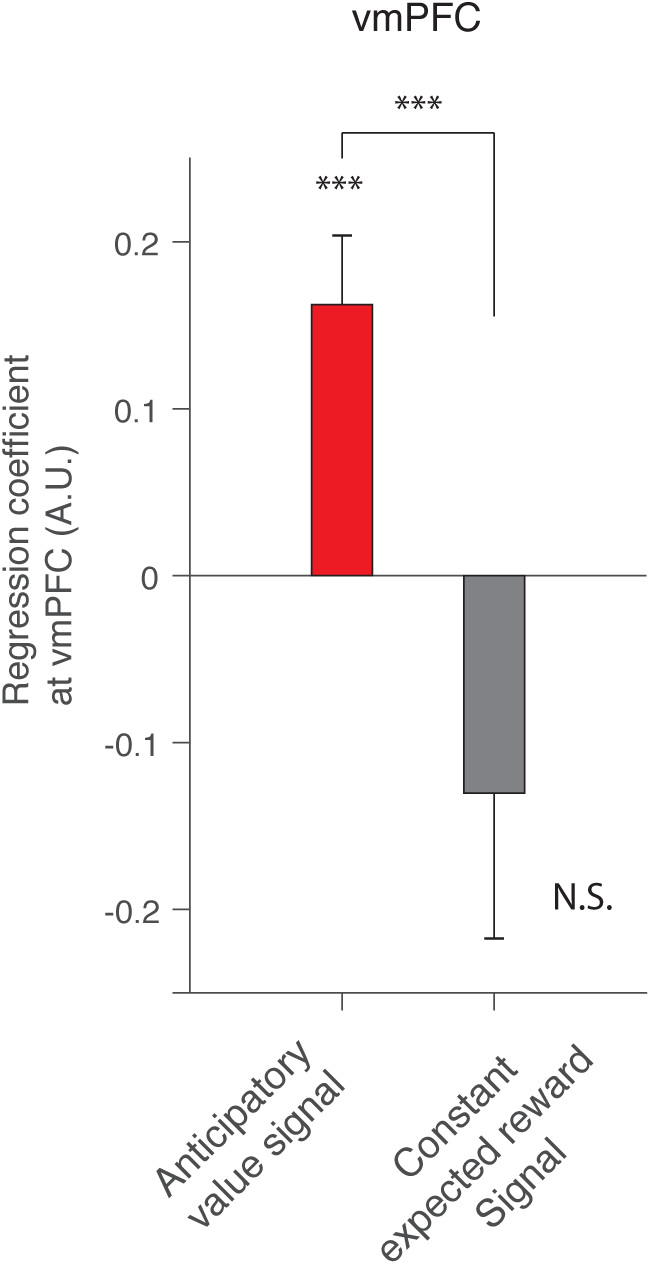
A control, confirmatory, analysis shows that the vmPFC is more strongly correlated with our model’s anticipatory utility signal than a standard expected reward value signal, defined by a boxcar regressor modulated by the probability of reward. In the GLM analysis with both regressors, average regression weights in the vmPFC cluster for the anticipatory value signal was significantly greater than the coefficients to the expected value signal (*p* < 0.001, permutation test). The average regression weights in the vmPFC cluster were significantly larger than zero for our model’s predicted signal (*p* < 0.001 t-test, *t*_38_ = 3.93), but not significantly different from zero for the expected value signal. The error bars indicate the mean and SEM. Note that this is a confirmatory analysis.

**Figure S11:**
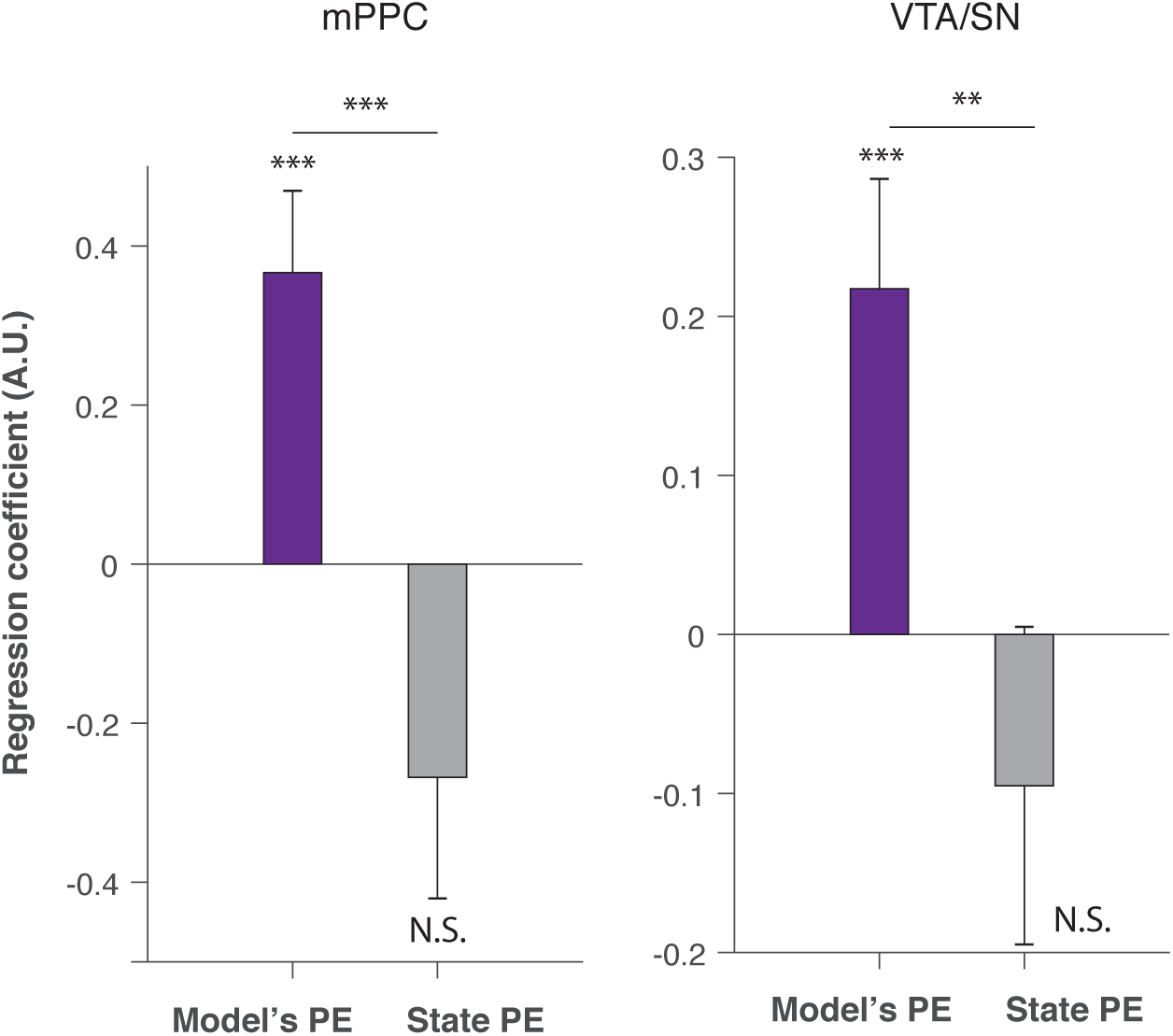
Our illustrative, confirmatory, analysis shows that BOLD signals in both the mPPC and the VTA/SN positively correlated with the model’s anticipation-reward prediction error signal (aRPE), but not with a simpler, so-called state prediction error signal (1 − *p*_reward_ when reward predictive cue was presented, |0 −*p*_reward_| when no-reward predictive cue was presented). The two regressors are included to the same GLM analysis. The differences between the average regression coefficients in the mPPC and in the VTA/SN were significant in the mPPC (*p* < 0.001, a standard permutation test in which we permuted the average regression coefficients), and the VTA/SN (*p* < 0.001 permutation test). The average correlation with the model’s aRPE signal was significant both in the mPPC and the VTA/SN (*p* < 0.001 for the mPPC and the VTA/SN; t-test *t*_38_ = 3.56 and *t*_38_ = 3.15). The three stars indicate *p* < 0.001, and two stars indicate *p* < 0.01.

**Figure S12:**
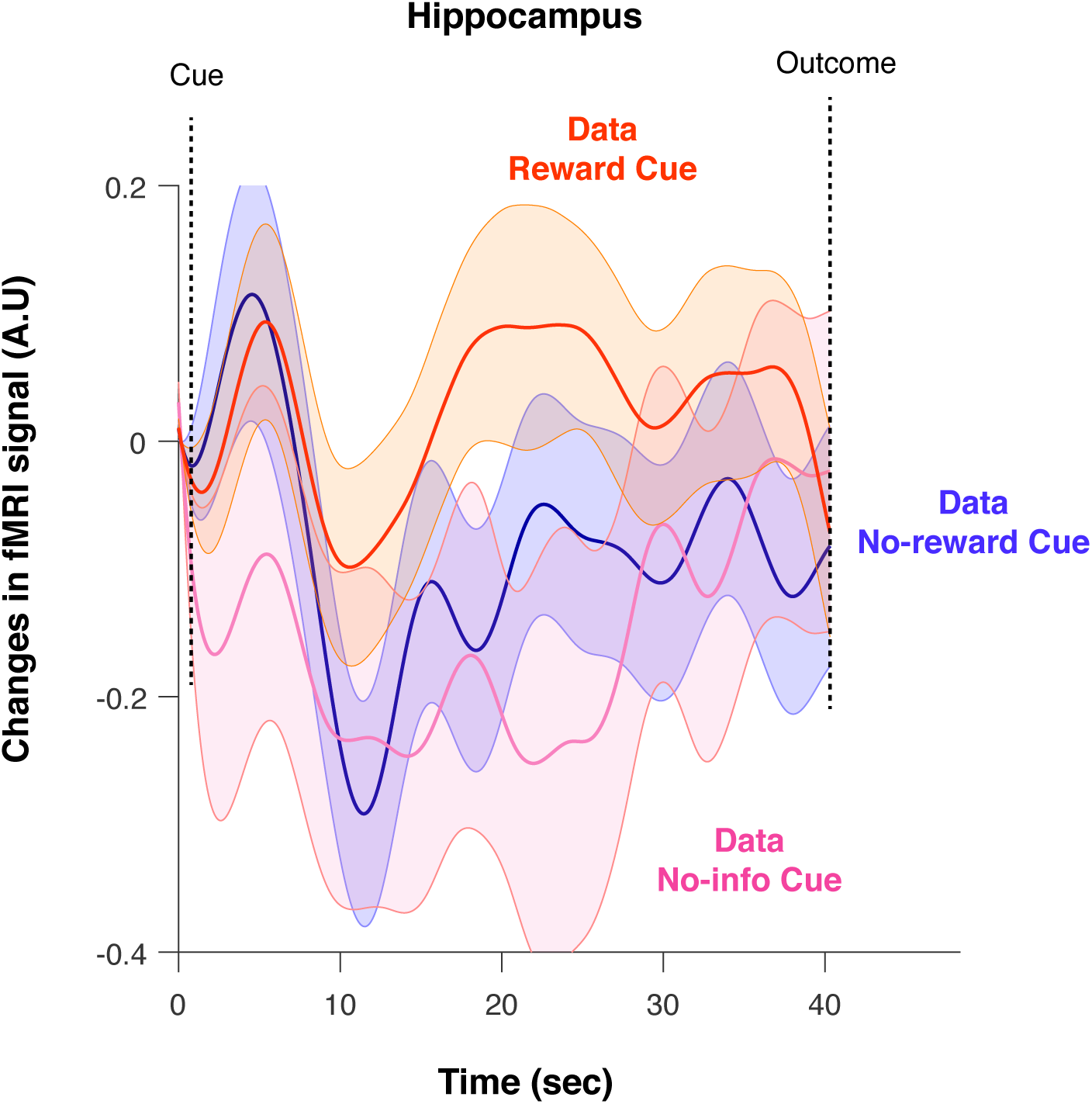
The temporal dynamics of the fMRI signal in hippocampus during anticipatory periods. Changes in activity averaged over participants after receiving a reward predictive cue (orange), after receiving a no-information cue (magenta), and after receiving a no-reward predictive cue (blue) are shown. The error bar indicates the SEM.

**Figure S13:**
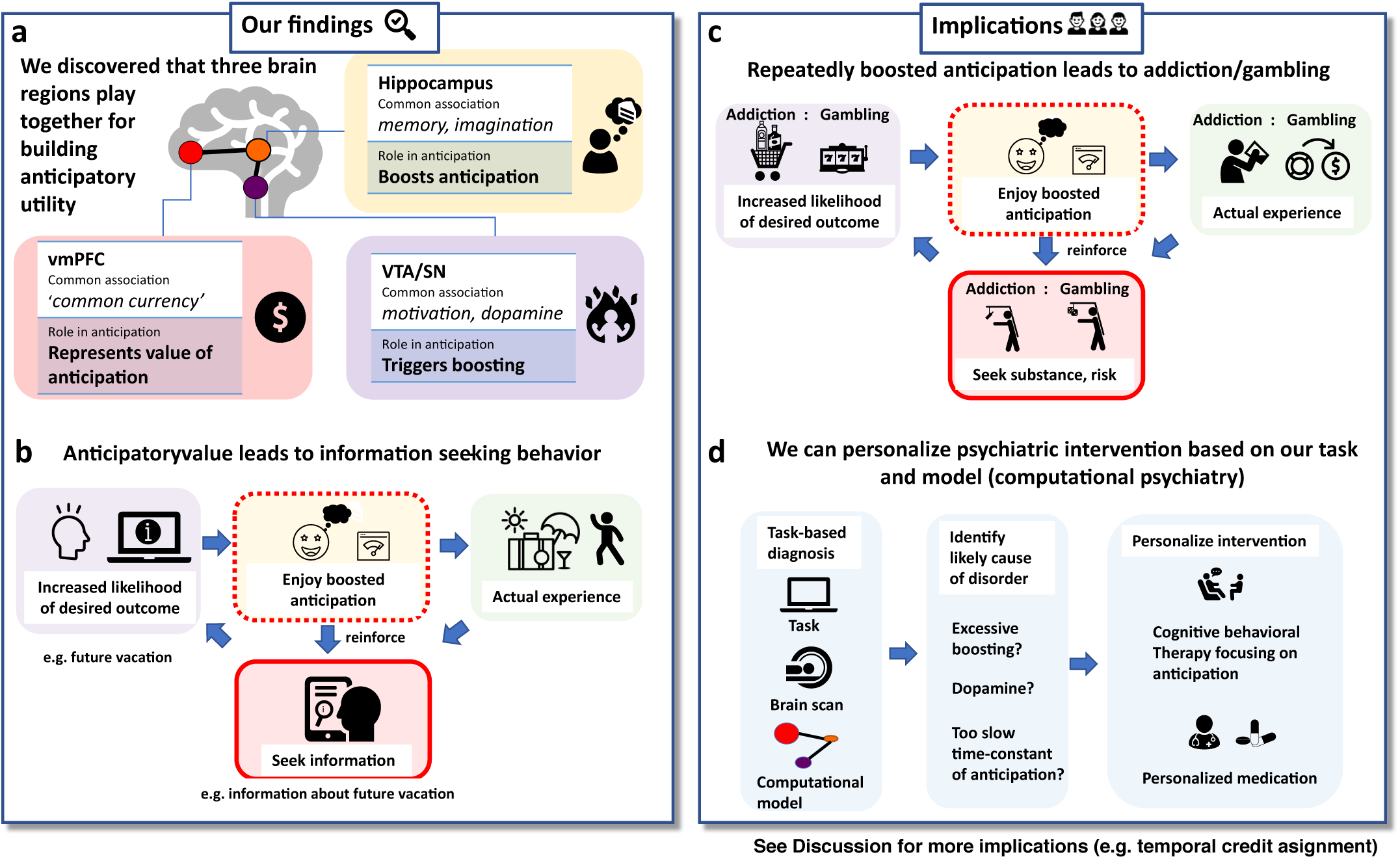
Summary of our findings and implications. (**a**,**b**). Our primary findings. (**a**). We uncovered three distinctive regions that play together to compute anticipatory value in the brain. The vmPFC, often described as brain’s common currency region, represented the value of anticipation. The VTA/SN, regions associated with dopamine and motivation, triggers the boosting of anticipation when the likelihood of desired outcome increases. The hippocampus, a strong associate of memory and imagination, realized the boosting of anticipation. (**b**). We showed that anticipatory value can drive information-seeking behavior. Advance information can increase the likelihood of a desired outcome, which in turn boosts the value of anticipation. As a result, people can feel enhanced value from anticipation after receiving advance information. Therefore people seek advance information of their desired outcomes (information-seeking, or observing), as we confirmed in our current and past experiments.^20^ (**c**,**d**). Implication of our study. (**c**). Over-boosted anticipation could lead to addiction and gambling. Purchasing alcohol or seeing ‘7-7-7’ in a slot machine increases the likelihood of receiving desired outcome (e.g. drinking alcohol, receiving money from gambling). This can boost the anticipatory value. By repeating this many times, the subjective value of alcohol or gambling can also be boosted over and over, leading to pathological seeking for substance (addiction) and risk (gambling). Note that our computational model predicts that this over-boosting can happen only to individuals with particular set of parameter values (e.g. strong boosting and weak discounting). (**d**). Our study can help to design personalized psychiatric interventions (computational psychiatry). Subjects perform a behavioral task in a MRI scanner and we fit our computational model to the behavior. We can identify likely causes of psychiatric disorders (e.g. addiction), by the subject’s parameters estimated by our computational model and brain data. This will help design personalized psychiatric intervention, for example cognitive behavioral therapy focusing on aspects of anticipation, as well as medication targeted to specific neuromodulators (e.g. dopamine). Please see the Discussion section for further details and other implications of our study.

